# PaleoRigor: a bounded research agent with a reproducible evidence trail for paleomicrobiome data processing

**DOI:** 10.64898/2026.09.30.755547

**Authors:** Tianao Liu, Haohong Zhang, Yiyang Jia, Kang Ning

## Abstract

**Background:** Ancient microbial evidence informs research on past health, diet, and environments, but its scarcity and susceptibility to contamination demand care in selecting and processing data. Language models can turn a research request into an analysis plan, but an unconstrained planner may reference files that do not exist, apply operations a format forbids, or answer a question the data cannot settle. A planner used on such evidence therefore needs to be bounded, and its decisions recoverable afterwards.

**Results:** We developed PaleoRigor around three commitments. Agent: a language model plans the analysis. Constraint: a validator checks every file reference, format, and step dependency before anything runs, execution is restricted to predefined local modules, and a planning contract directs the model to refuse out-of-scope requests with a named reason code rather than answer them. Reproducibility: each run is exported with its plan, intermediate files, checksums, and decisions. In a preregistered, frozen held-out evaluation, PaleoRigor passed 23 of 24 runs (95.8%; Wilson 95% confidence interval, 79.8–99.3%): 11 of 12 supported runs and all 12 prespecified boundary decisions. An ablation that removed this planning contract from the same model passed 19 of 24 (79.2%). Repeating the frozen design with a second model configuration yielded 24 of 24 successes for PaleoRigor and 19 of 24 for the ablated arm. Pooled post hoc across both configurations, PaleoRigor passed 47 of 48 runs (97.9%), compared with 38 of 48 (79.2%) for the ablated arms. All 96 run labels were recomputed from retained records and matched. Dataset-level tests matched read counts and compressed-file sizes for six public sequencing records and documented the removal of 114 duplicate sequence entries. A retraction-associated audit illustrated how expert review limited conclusions to file- level evidence.

**Conclusions:** For paleomicrobiome data, pairing a language-model agent with explicit constraints and a reproducible record makes the agent’s proposed analysis reviewable before its outputs support a biological claim. PaleoRigor delivers checked inputs, a recorded transformation history, and a named reason for each request it refuses, and runs as a local desktop application. It prepares data for specialist analysis; ancient-DNA authentication and biological interpretation remain with the paleobiologist.

**Key points:**

- **Agent:** a language model turns a research request into an explicit, ordered workflow, which a paleobiologist reviews and approves before anything runs. In the frozen held-out evaluation, which ran without that review step, 11 of 12 supported runs passed every planning, validation, and execution check.
- **Constraint:** a validator, a fixed registry of local skills, and encoded refusal rules bound what the agent may do. The refusal rules direct the planner to decline, with a named reason code, requests that cross a limit (for example, establishing authenticity or absence of contamination from quality- control output). In the frozen v5 round, all 12 boundary decisions were correct, compared with 10 of 12 when the planning contract was removed.
- **Reproducibility:** every run links the request to its inputs, planned operations, intermediate files, checksums and, for refused requests, the blocked decision; all 96 run labels from the two frozen model configurations were recomputed from these retained records and matched.

**Graphical abstract:** 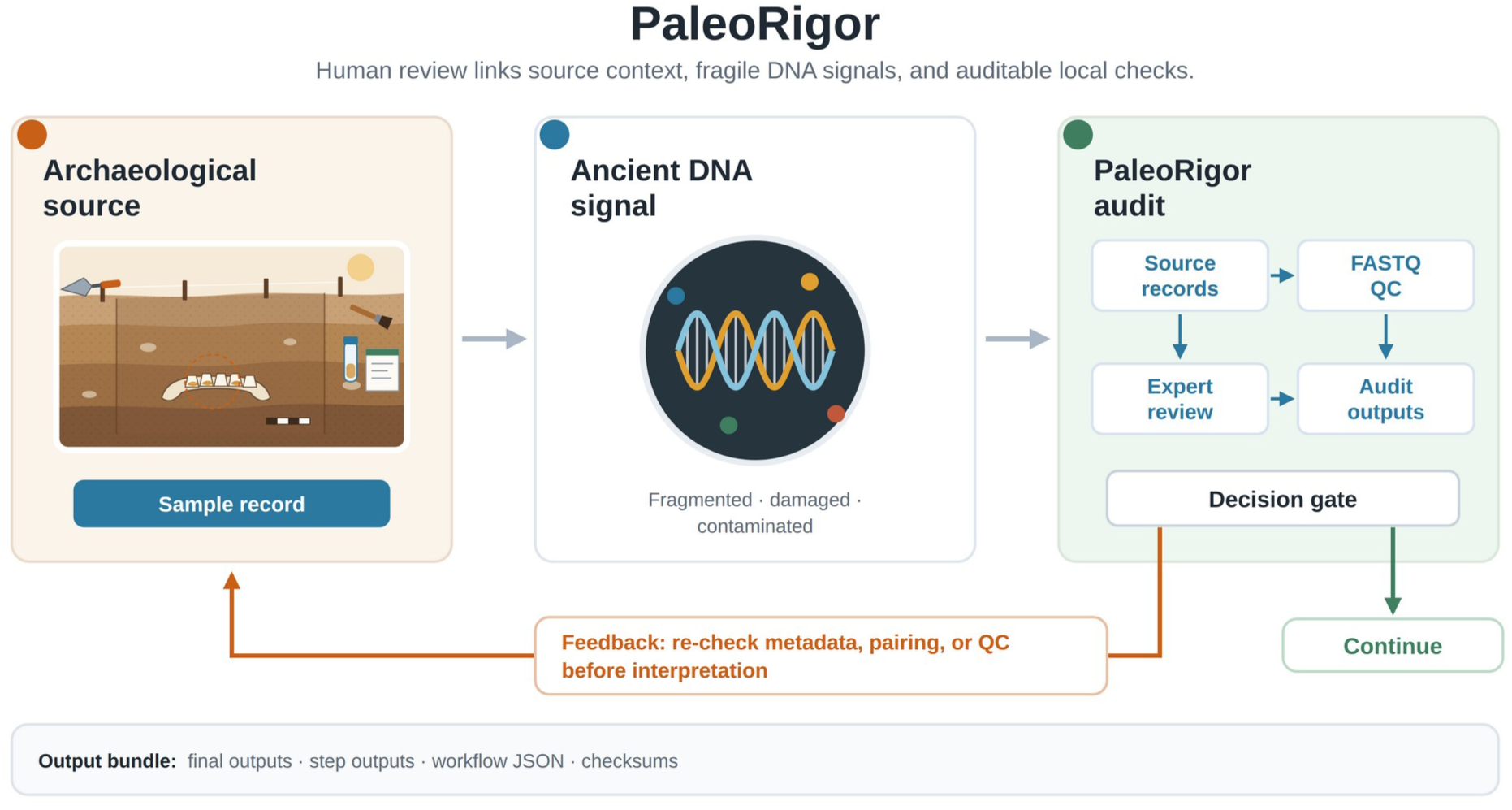

Graphical abstract. PaleoRigor connects archaeological source material, fragile ancient biomolecular evidence, and reviewable local analysis. All elements, including the schematic excavation scene, were drawn by the authors as vector graphics; the graphical abstract contains no photographs. The graphical abstract shows the paleobiological use case, while Figure 1 shows the expert review and local execution steps. In the figure, the decision gate marks the expert’s choice between continuing and re-checking the source record.

## Background

Paleomicrobiome and ancient-human microbiome studies examine archaeological material to investigate past health, diet, and microbial exposure [1–5]. Sequencing data are often deposited in the Sequence Read Archive (SRA) and European Nucleotide Archive (ENA) [6, 7], and community curation has standardized the sample-level metadata that accompanies published ancient metagenomic records [8]. Access to these data is only the starting point for analysis. Researchers must select the intended files, check their local copies, choose appropriate operations, and document subsequent changes. Errors at this stage can persist through an otherwise successful analysis. A traceable connection between the research question, repository record, and local workflow is therefore essential.

These requirements are particularly consequential in ancient biomolecular research, where DNA is often scarce, fragmented, damaged, and contaminated. Authenticity depends on several sources of evidence, including sample history, molecular damage, contamination controls, and archaeological context [3, 9–13], and the field has published explicit protocols for authenticating ancient DNA recovered from metagenomic data [14]. Mapping, adapter removal, and damage analysis assess the files supplied to them [15–18]; they cannot compensate for an incorrect starting file. Source verification therefore supports, but does not replace, biological authentication.

Standardized workflows can reduce the manual file selection and handling on which such checks depend, and make complex analyses easier to review. Generic workflow engines support reproducibility once users have specified inputs and operations [19–21]. The ancient-metagenomics community has built domain pipelines on them: aMeta implements profiling and authentication as a Snakemake workflow [22], nf-core/eager implements genome reconstruction in Nextflow [23], HOPS couples pathogen screening to automated authentication [24], and more recent workflows continue to optimize the profiling step itself [25]. In these pipelines the user has already chosen the inputs and operations, and deciding whether those choices suit the research question still requires expertise. Language-model agents can now translate a request into such choices directly [26], which removes the need to write the plan by hand but introduces a new failure mode: the plan is generated rather than authored, and may reference files that do not exist, apply operations a format does not permit, or assert conclusions the data cannot support. Neither an unconstrained agent nor a workflow engine records why each step of the executed plan was chosen, so such an error is hard to trace afterwards. What is therefore missing is a mechanism that states a proposed analysis explicitly, constrains what it may do before it runs, and preserves the reasoning and evidence afterwards, while leaving responsibility for the biological conclusions with the researcher.

We developed PaleoRigor as a bounded research agent for this purpose, built on three commitments. First, it is an agent: its planner converts a research request into a proposed workflow, so that the researcher reviews a plan rather than writing one. Second, the agent acts under constraint: its validator checks file references, formats, and compatibility between steps; each step must call a skill, a predefined local analysis module that the user can inspect; and refusal rules direct the agent to decline, with a named reason code, requests that cross an encoded limit on interpretation or on required inputs. The agent cannot write or execute code outside that skill registry, so the space of possible actions is fixed before the model is called. Third, each run can be reconstructed from its record: after approval, the intermediate and final outputs of the selected skills are saved, and a report links those outputs to the workflow and its execution records, so that the outcome of a run can be recomputed without calling the model again. This division allows paleobiologists to review the proposed analysis and assess its results without delegating scientific judgment to the planning model (Figure 1; Table 1).

**Table 1.**
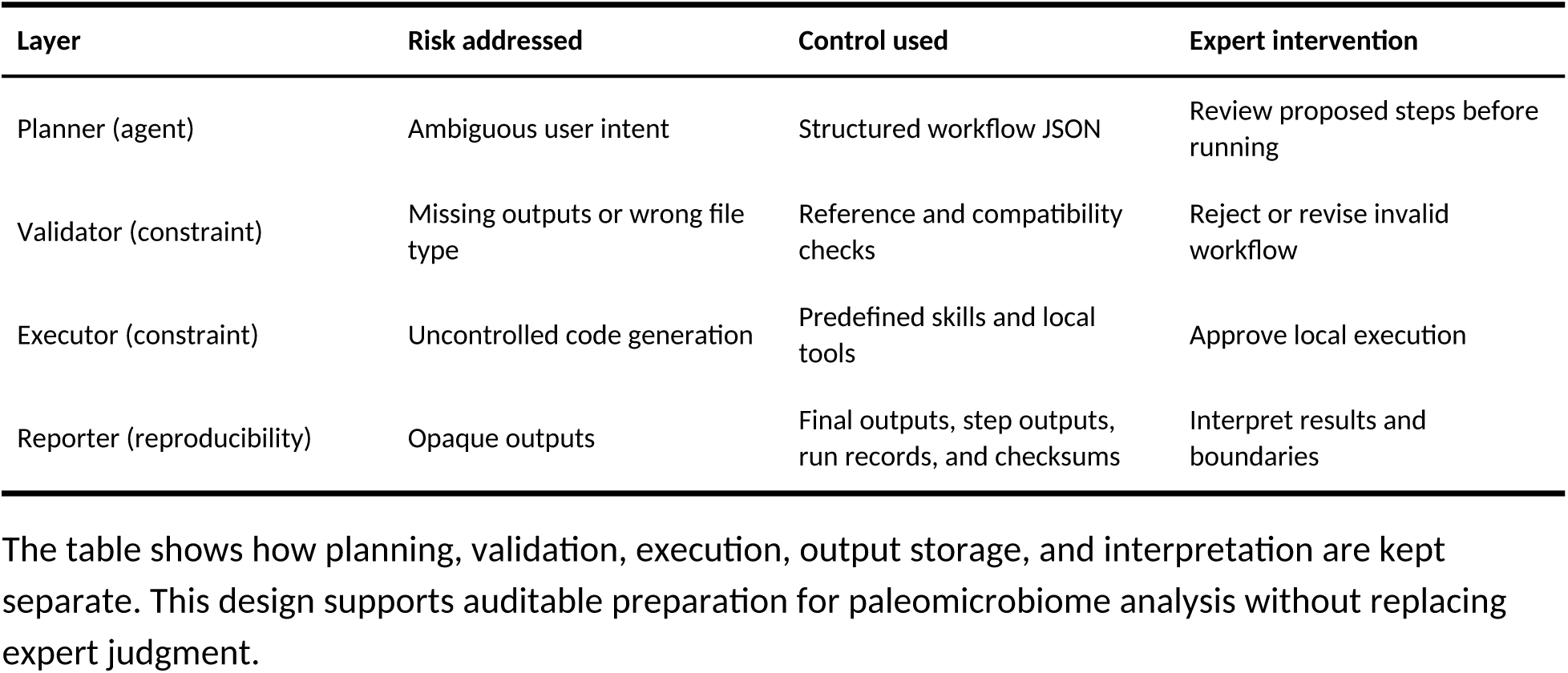
PaleoRigor control layers and expert checkpoints.

We evaluated each commitment. A frozen held-out benchmark tested the agent and its constraints together: whether supported requests produced the required outputs and whether unsuitable requests were blocked for the correct reason, compared with an ablation that removed the system’s own planning contract (the planning and refusal rules supplied to the model). Reproducibility was tested by recomputing every benchmark label from the retained records, and two dataset-level cases examined what those records make checkable: agreement between local sequencing files and public records, and a recorded table transformation. A third case, an expert-defined audit of a retraction-associated record, examined the constraints on interpretation from the expert’s side. Together, these tests assess preparation for specialist analysis, rather than ancient-DNA authentication or biological reconstruction. The project website, https://verture-liu.github.io/PaleoRigor/, explains how PaleoRigor is used step by step, links to the desktop applications, and presents every held-out benchmark run for inspection [27].

## Results

### Agent, constraint, and reproducibility: control layers and test criteria

PaleoRigor implements its three commitments as four control layers, each addressing one of four risks that precede paleobiological interpretation (Table 1): ambiguous user intent, missing outputs or wrong file types, uncontrolled code generation, and opaque outputs. The planner is the agent: it states the request as an explicit ordered workflow. The validator and executor impose the constraints: the validator checks file references and format compatibility, and the executor runs only predefined skills; refusal rules in the planning contract add a further constraint, described below. The reporter provides reproducibility: it stores final outputs separately from the records that explain them (Figure 1).

Expert review is placed at three points in that sequence: when the request is framed, before execution, and when the retained evidence is interpreted (Additional file 1: Table S1). Table 1 sets out which layer addresses which risk, and where the paleobiologist rather than the planning model remains responsible for the decision.

The refusal rules cover a risk that checks on the plan cannot remove, because it lies in the request itself: a request that the supplied data cannot support, either because it asks for a conclusion the evidence cannot establish or because a required input is missing or of the wrong kind. The limits that matter most in ancient biomolecular work are stated to the planner as refusal rules rather than left to the user to remember. For conclusions, quality-control output cannot establish that reads are ancient or free of contamination, so the planner is directed to refuse a request to demonstrate either rather than answer it. For inputs, damage analysis requires aligned data and an explicitly named reference [15, 18], so it is refused when those prerequisites are absent. Read alignment and host removal are refused when no reference genome or existing index is named, instead of defaulting to one. A request whose file format contradicts the operation is refused rather than silently replaced by a different analysis, and a request that names a paired-end mate that was not supplied is refused rather than completed with an invented or reused file. Each refusal returns one of four named reason codes, which follow these two kinds of limit: unsupported_scientific_claim for conclusions, and missing_prerequisite, file_type_mismatch, and missing_mate for inputs; the rules are reproduced verbatim in Additional file 2.

To test whether the control layers and refusal rules work as intended, we defined six criteria (Table 2) covering workflow validity, file compatibility, source agreement, traceability, software dependencies, and repeated-run workflow success with boundary control. These criteria were fixed before the frozen held-out evaluation reported below. A supported run passed only if it met every applicable planning, validation, and execution requirement; a boundary run passed only if it stopped with the prespecified reason. Scoring was therefore strict, and we report the proportion of runs passing on this basis as strict success.

**Table 2.** Engineering criteria used to assess PaleoRigor.

| **Evaluation target** | **Metric or rule** | **Interpretation** |
|---|---|---|
| Workflow validity | All step inputs must reference uploaded files or earlier outputs | Prevents hallucinated pipeline references |
| File compatibility | Input format must match skill requirements | Prevents applying FASTQ operations to CSV files |
| Source agreement | Δreads = 0 and Rbytes = 1 when public metadata are available | Confirms local files match repository records at the metadata level |
| Traceability | Final outputs plus step outputs and run records | Allows later inspection by users or supervisors |
| Dependency behavior | Missing tools are reported instead of auto-installed | Keeps execution transparent on user machines |
| Repeated-run workflow success and boundary control | Supported tasks must pass all tests; incompatible or overreaching requests must stop before unsafe execution | Separates repeatable workflows from safe scientific boundaries |

Each criterion addresses a type of error seen during development and states how the system should prevent or expose it before biological interpretation. Δreads is the local read count minus the public read count. Rbytes is the local compressed FASTQ size divided by the public fastq\_bytes value. CSV, comma-separated values.

### Agent and constraint: a frozen held-out evaluation

The frozen evaluation was preceded by two development-stage qualification rounds (Figure 2A). Each comprised four supported and four boundary scenarios, repeated three times in each of two arms (PaleoRigor and an ablated arm, described below), and PaleoRigor passed 18 of 24 runs in v3 and 18 of 24 in v4 (75.0% in each round), compared with 18 of 24 and 21 of 24 for the ablated arm; these rounds showed no advantage for PaleoRigor. Failures arose when declared output names did not match skill outputs, chart-name variants were rejected, and sample-sheet validation required unavailable FASTQ files. These rounds informed corrections to output declarations and planning rules. Because they shaped the system, they cannot serve as independent validation of it, and their failures are retained in the reported record.

**Figure 1.**
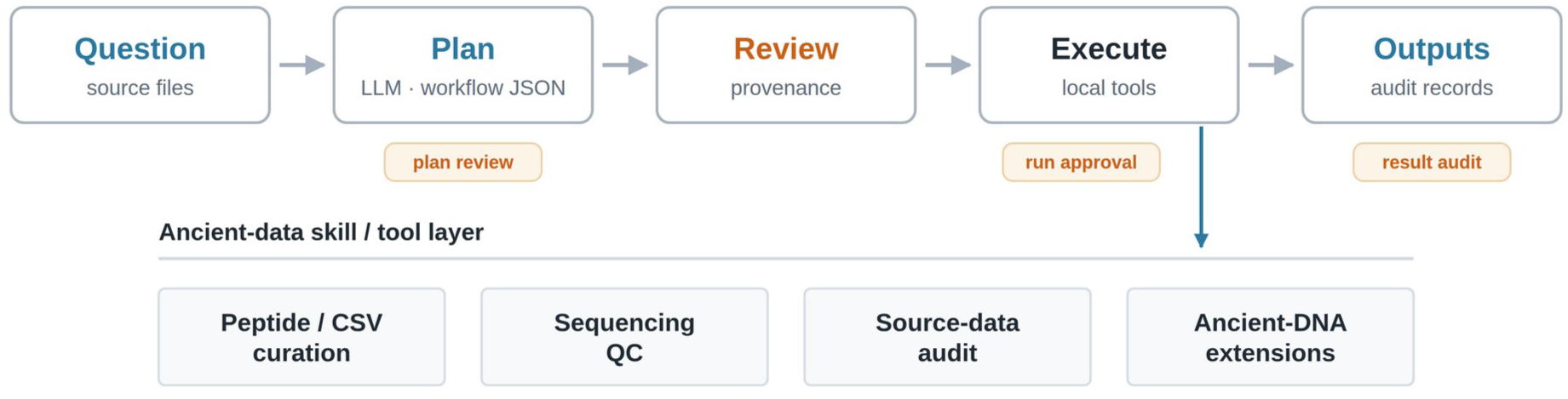
Human-in-the-loop design of PaleoRigor. A paleobiologist starts with a research question and available source material, such as metadata tables, sequencing files, or repository records. PaleoRigor drafts a structured workflow. Experts then review the source, file organization, task fit, and parameters before local execution; the orange tags mark the three expert checkpoints (plan review, run approval, and result audit). Approved skills perform limited tasks such as table cleaning, sequencing quality control, source-record comparison, and ancient-DNA extensions (alignment and host removal). Final outputs are stored separately from step outputs and run records so that readers can inspect both the result and how it was produced. In the figure, LLM means large language model, QC means quality control, and workflow JSON (JavaScript Object Notation) means a machine-readable text record of the proposed plan. In the terms used in the text, drafting the workflow is the agent’s role; reference and format checks and execution limited to approved skills are the constraints; and storing final outputs separately from step outputs and run records provides reproducibility (Table 1).

**Figure 2.**
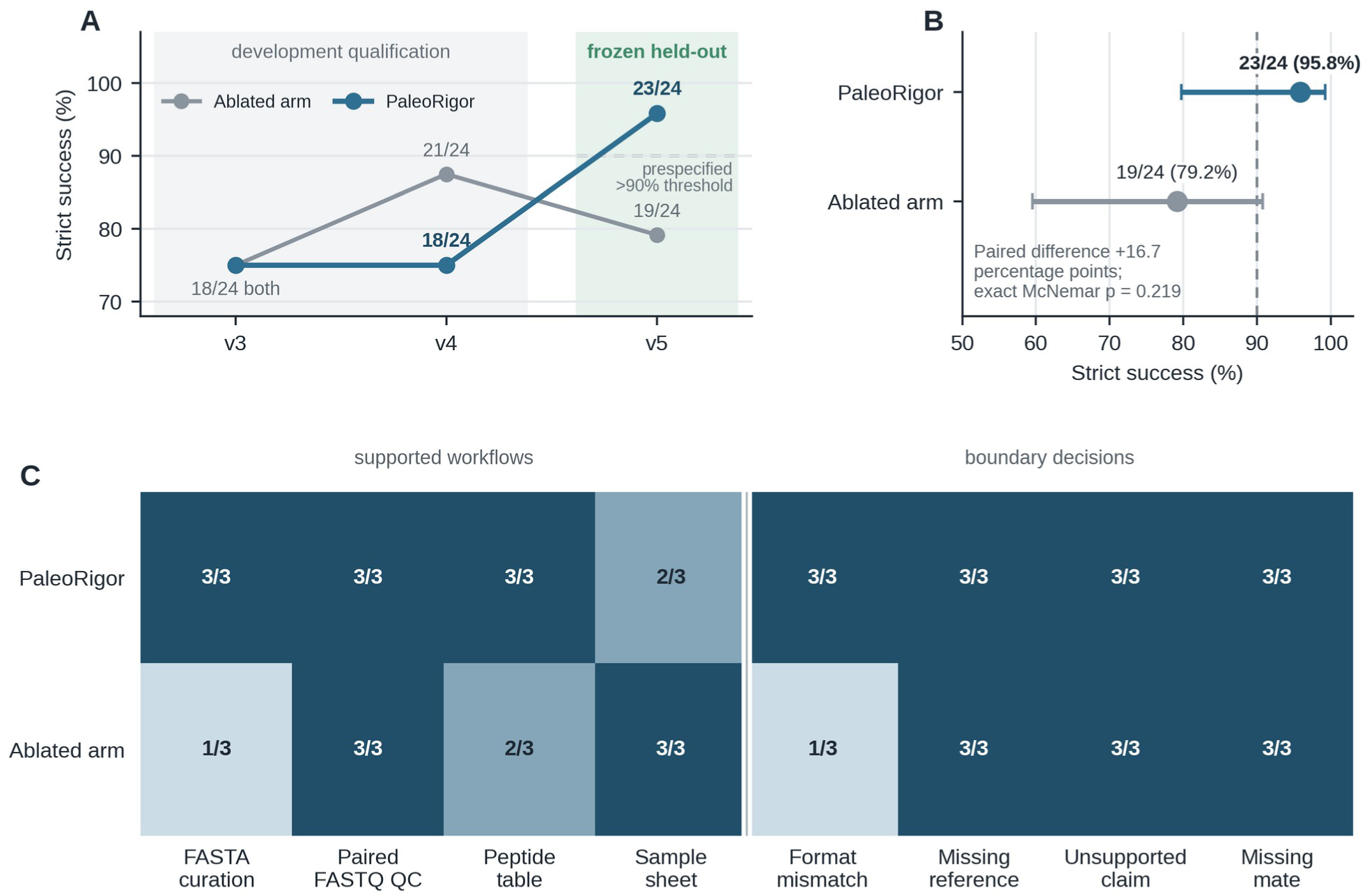
Versioned engineering benchmark and frozen held-out evaluation. A, Strict run-level success across v3, v4, and v5. Rounds v3 and v4 were retained as development-stage qualification rounds; each contained 24 PaleoRigor and 24 ablated- arm runs. The frozen v5 round used new files, identifiers, values, and wording. The dashed line marks the prespecified v5 threshold of more than 90%, operationalized as at least 22 of 24 successful PaleoRigor runs. B, Frozen v5 strict success with deepseek-v4-flash for PaleoRigor and for the ablated arm, in which the PaleoRigor planning contract was removed from the same model. The dashed line marks the same threshold. Points show observed proportions and horizontal bars show two- sided Wilson 95% confidence intervals. The paired difference was 16.7 percentage points; the exact two-sided McNemar p value was 0.219. C, Successes by scenario in v5 with deepseek-v4-flash. Supported workflows comprised FASTA curation, paired FASTQ quality control, peptide-table processing, and sample-sheet preparation. Boundary decisions tested a file- format mismatch, a missing reference, an unsupported authenticity or contamination claim, and a missing paired-end mate. Each cell reports successful runs out of three repeats with identical inputs; repeats are not independent samples.

To test the agent and its constraints together, we then froze the manifest, inputs, prompts, scoring rules, threshold, and code before the first call and ran v5, the last of three versioned benchmark rounds (Figure 2A), on held-out material: new files, identifiers, values, and request wording. Four supported scenarios tested the agent, asking whether it planned a workflow that passed validation and executed; four boundary scenarios (a file-format mismatch, a missing reference, an unsupported authenticity or contamination claim, and a missing paired-end mate) tested the constraints, asking whether it refused with the prespecified reason code. Each scenario was repeated three times per arm, producing 24 runs for PaleoRigor and 24 for an ablated arm, with deepseek-v4-flash as the planning model.

That arm was an ablation of the PaleoRigor planning contract, not a trial against an external tool. Both arms used the same model, task files, call-order protocol, validator, execution path, and scoring rules, and received the same workflow schema, blocked-decision schema, uploaded-file summaries, and skill registry as a byte-identical shared prompt block. The arms differed only in the planning instructions preceding that block: the ablated arm received a 247-character, four-sentence planner instruction, and the PaleoRigor arm the full 40,347-character planning contract, whose six refusal rules name the four boundary reason codes (Methods). The boundary comparison therefore measures whether an explicitly encoded rule is applied consistently; it does not measure whether a planner identifies a boundary it was not told about. Both system prompts are reproduced verbatim in Additional file 2.

PaleoRigor passed 23 of 24 runs (95.8%; Wilson 95% confidence interval, 79.8–99.3%): 11 of 12 supported runs and 12 of 12 boundary decisions. This exceeded the prespecified threshold of at least 22 of 24. The ablated arm passed 19 of 24 (79.2%; 59.5–90.8%): 9 of 12 supported runs and 10 of 12 boundary decisions (Figure 2B). The paired difference was 16.7 percentage points; five runs passed only with PaleoRigor and one only with the ablated arm (exact two-sided McNemar p = 0.219), so the comparison did not establish statistical superiority. All 48 outcomes were recomputed from the retained records, and all 48 matched the stored labels, so the scoring of every run can be reproduced from its retained records without calling the model again.

Scenario-level results locate the remaining errors (Figure 2C); each run can also be inspected, scenario by scenario and for both model configurations, in the benchmark explorer on the project website (https://verture-liu.github.io/PaleoRigor/explorer.html). PaleoRigor passed 3 of 3 runs for FASTA curation, paired FASTQ quality control, peptide-table processing, and each of the four boundary scenarios, and 2 of 3 runs for sample-sheet preparation. In the failed run the planner omitted the .tsv suffix from an uploaded-file reference, which the validator rejected before execution: the constraints caught a planning error before it could produce an output. The ablated arm passed 1 of 3 runs for both FASTA curation and file-format mismatch decisions, and 2 of 3 peptide-table runs.

To check whether these results depended on a single planning model, we repeated the frozen design with a second model configuration from the same provider, deepseek-v4-pro, changing only the requested model (Methods). PaleoRigor passed 24 of 24 runs (100%; Wilson 95% confidence interval, 86.2–100%) and the ablated arm 19 of 24 (79.2%; 59.5–90.8%; 8 of 12 supported runs and 11 of 12 boundary decisions), a paired difference of 20.8 percentage points with all five discordant pairs favoring PaleoRigor (exact two-sided McNemar p = 0.0625). This met the preregistered criterion of at least 22 PaleoRigor successes and more successes than the ablated arm. All 48 second-model labels were likewise recomputed from the retained records and matched.

Both configurations therefore met their prespecified engineering criteria on the evaluated system and task set. Across both, PaleoRigor passed 47 of 48 runs (97.9%) and the ablated arms 38 of 48 (79.2%); this pooled summary was not prespecified, and the two configurations were not used to rank models.

The three dataset-level cases below turn from benchmark scenarios to individual datasets (Table 3) and the records retained for them. Two extend the reproducibility test beyond the label recomputation reported above: whether the local sequencing files a run starts from are the intended public records (Figure 3), and whether a table transformation can be checked exactly from its run record (Figure 4). The third examines the constraints on interpretation from the expert’s side, using a record whose published interpretation was retracted; here the limit was set by the expert rather than encoded in the system, and Figure 5 shows where file-level evidence stops. In Tables S2–S4 of Additional file 1, they appear as Case 1, sequencing source checks; Case 2, peptide-table curation; and Case 3, retraction-associated audit.

**Figure 3.**
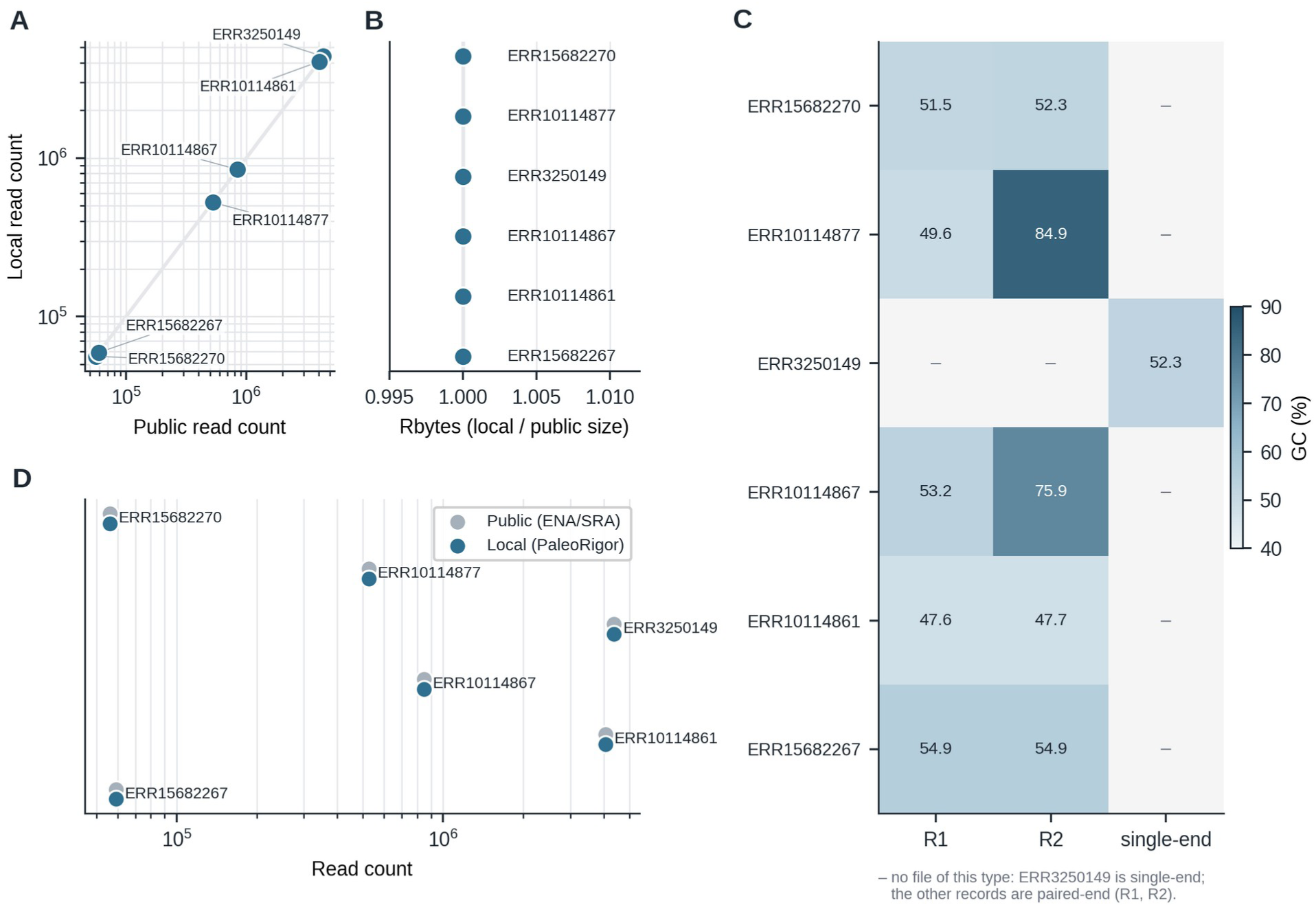
Agreement between local sequencing files and public records. A, Local read counts compared with ENA/SRA read counts. B, Ratio of local compressed-file size to public fastq_bytes (Rbytes). For paired-end records, local counts and sizes are summed over both mates. Δreads is local reads minus public reads. Rbytes is local compressed bytes divided by public fastq_bytes. C, GC content for R1, R2, or single-end files; dashes mark file types a record does not have. D, Public (ENA/SRA) and local (PaleoRigor) read counts for each record. Read counts and sizes alone cannot prove byte-level identity, which was confirmed separately with MD5 checksums (Methods); none of these checks establishes ancient-DNA authenticity or biological validity.

**Figure 4.**
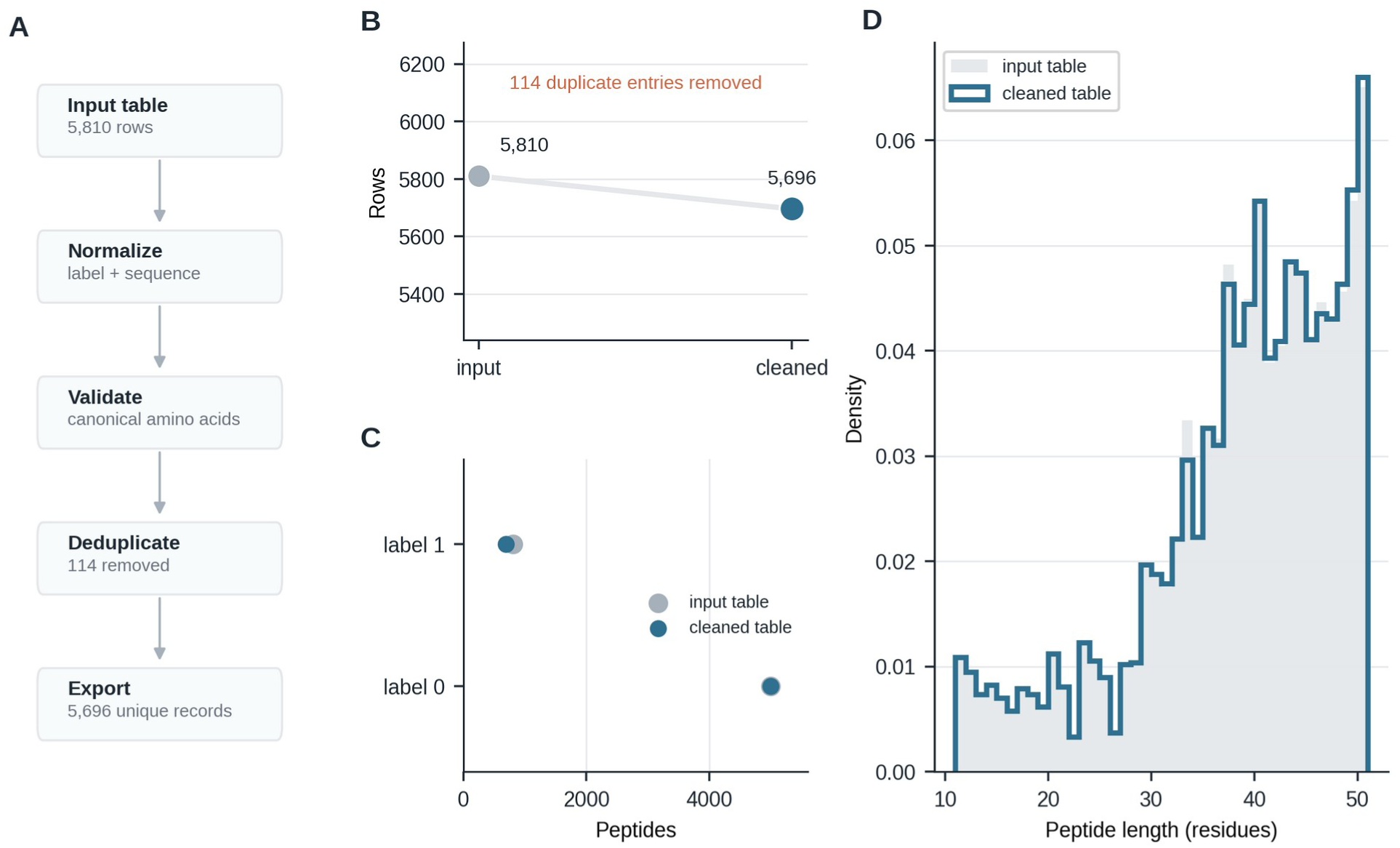
Peptide-table curation. A, Steps used to process the peptide table. B, Row count before and after duplicate removal, from 5,810 input rows to 5,696 unique records. C, Counts for the two peptide labels before and after cleaning. D, Peptide-length distribution (density) before and after cleaning, measured as amino-acid residues per sequence.

**Figure 5.**
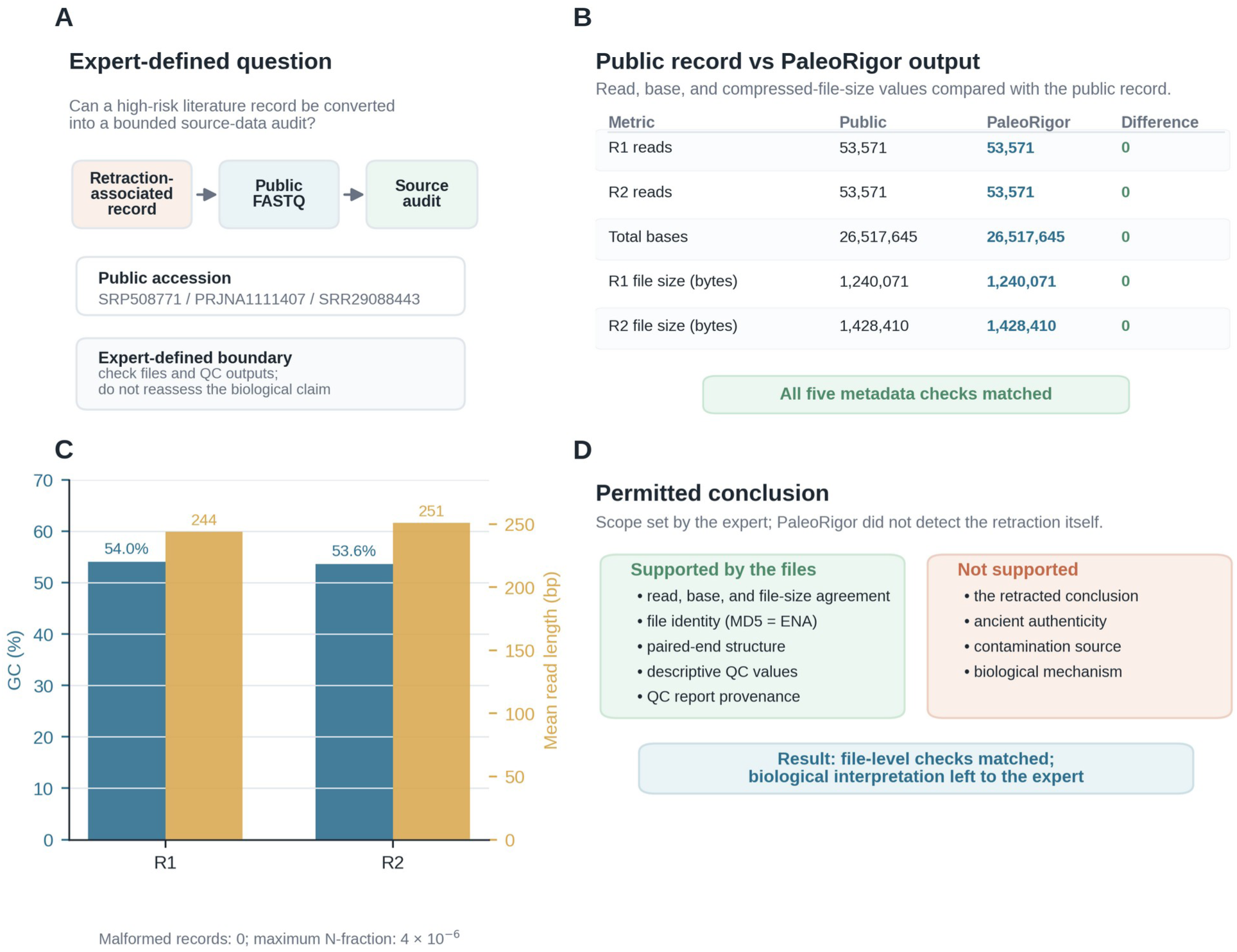
File-level audit of a retraction-associated FASTQ record. Retraction-associated refers to the linked publication context, not to an autonomous retraction decision by PaleoRigor. The modern clinical microbiome record was used as a negative-control test of interpretation boundaries, not as an ancient-microbiome dataset. A, The expert limited the question to a source audit. B, Public metadata and PaleoRigor outputs matched for R1 reads, R2 reads, total bases, and compressed- file sizes. R1 and R2 are paired-end mates. C, Descriptive quality-control results from the same files, including GC content, mean read length, malformed-record checks, and N-fraction. N-fraction is the proportion of bases recorded as N. D, The permitted conclusion was limited to file-level reproducibility, including file identity confirmed afterwards with MD5 checksums (Methods), and did not validate the retracted biological interpretation.

**Table 3.** Datasets and measured results used in the evaluation.

| Evaluation dataset | Input and original source | PaleoRigor task | Quantitative readout or expected boundary |
|---|---|---|---|
| Sequencing source checks | Six public sequencing records: ERR15682270, ERR10114877, ERR3250149, ERR10114867, ERR10114861, and ERR15682267. Original ENA accessions listed above [6, 7] | Confirm local files match intended ENA/SRA records before downstream analysis; generate FastQC, MultiQC, and SeqKit summaries. | PaleoRigor-derived read counts matched public metadata with Δreads = 0; local compressed-file ratios were Rbytes = 1; MD5 checksums matched ENA for all 11 files; GC content and mean read length were reported as quality-control metrics. |
| Peptide-table curation | Headerless local Validation.csv peptide table; 5,810 rows. Project repository [33] | Normalize columns, validate peptide sequences, remove duplicate sequences, compute summaries, and export a cleaned table. | 5,696 unique records exported; 114 duplicate sequence entries removed; label balance and peptide-length distribution retained as figure checks. |
| Retraction-associated audit | Paired FASTQ record SRR29088443 from SRP508771 / PRJNA1111407; 53,571 reads per mate and 26,517,645 total bases. Original SRA and BioProject accessions listed above [6] | Test whether a problematic literature-linked record can be converted into a source-data audit without validating the retracted biological claim. | R1 reads, R2 reads, total bases, R1 file size, and R2 file size matched the public record exactly; biological interpretation was treated as expert-gated. |
| Engineering benchmark | Versioned v3–v5 benchmark and a preregistered second-model check of the frozen v5 design; each used eight scenarios, three repetitions, and matched PaleoRigor and ablated arms (24 runs per arm). Project repository [33] | Test strict workflow success, local execution, input/scientific boundaries, and the effect of the PaleoRigor planning contract under fixed model settings. | PaleoRigor v3: 18/24; v4: 18/24; frozen v5 (deepseek-v4-flash): PaleoRigor 23/24 (95.8%) and the ablated arm 19/24 (79.2%). Boundary decisions in v5: 12/12. Second-model check (deepseek-v4-pro): PaleoRigor 24/24 (100%) and the ablated arm 19/24 (79.2%). |

The table lists each input, its public source, the PaleoRigor task, and the measured output or required boundary. Public accessions identify the source data. The reported values show what was reproduced without making a biological claim. GC, guanine–cytosine; MD5, Message-Digest algorithm 5.

### Reproducibility: agreement between local sequencing files and public records

This case tested what the retained record makes checkable at the input end: whether local sequencing files agreed with the intended repository records before ancient-DNA interpretation. Six public records related to ancient or paleomicrobiome research were examined. PaleoRigor checked file structure, ran FastQC, summarized reports with MultiQC, calculated SeqKit statistics, and compared local files with ENA/SRA metadata [6, 7, 28–30]. Run records retained the accession, library layout, and mate organization.

All six records matched public read counts exactly (Δreads = 0), and compressed-file sizes matched repository values (Rbytes = 1; Figure 3A,B). The Message-Digest algorithm 5 (MD5) checksums of all 11 analysed files also matched those published by ENA (Methods), so the analysed files were identical to the public ones. These checks confirm that each run started from the intended record and provide an initial check of source selection before downstream processing [15, 17]. Guanine–cytosine (GC) content and a per-record comparison of read counts are shown in Figure 3C,D. These values are descriptive: neither read length nor GC content establishes ancient-DNA authenticity or excludes contamination [3, 14].

### Reproducibility: a recorded transformation of a data table

This case tested the reproducibility of a data change: whether the saved run record alone could explain a change to a data table. We used a peptide table because duplicate removal provides a small, exactly checkable transformation. PaleoRigor standardized columns, checked sequence syntax, removed 114 duplicate sequence entries from 5,810 input rows, and exported 5,696 unique records (Figure 4A,B).

The run record attributed the removal of the 114 duplicate sequence entries to the deduplication step. Class counts and peptide-length distributions provided additional checks for unintended changes (Figure 4C,D). This documented transformation satisfied the traceability criterion in Table 2.

### Constraint: expert-defined limits on interpretation for a retraction-associated record

This case examined how expert review could limit conclusions when technically reproducible files were associated with a retracted publication. We selected SRR29088443 from SRP508771 as such a record: the accession itself was not marked as retracted, but the SRA study title and submitting center matched the publication record [31, 32]. Because these data originated from a modern clinical microbiome study, the case served as a negative control for interpretation boundaries rather than as an ancient- microbiome analysis. The published interpretation had been retracted, so the expert restricted the workflow to file-level checks and excluded reassessment of the retracted biological claim (Figure 5A).

Within the approved scope, PaleoRigor reproduced the repository values. Each FASTQ mate contained 53,571 reads, with 26,517,645 bases across both files, both compressed-file sizes matched the public record, and the MD5 checksums of both files matched those published by ENA (Methods). Quality- control summaries reported mean read lengths of 244 and 251 base pairs (bp) and GC contents of 54.0% and 53.6% for R1 and R2, respectively (Figure 5B,C).

Figure 5D makes the division of responsibility explicit. The expert supplied the retraction context and defined the permitted analysis; PaleoRigor performed the file checks and retained their outputs. The software did not independently detect or adjudicate the retraction. The case illustrates how expert- defined scope can prevent agreement with repository metadata from being presented as biological validation.

Table 4 brings together the measured outcomes of the benchmark and the three dataset-level cases and sets them alongside common properties of existing workflow practice; no external workflow tool was measured.

**Table 4.** Operational comparison of PaleoRigor with existing workflow practices.

| Commitment and tested property | Measured PaleoRigor outcome | Contrast with common practice (not measured) |
|---|---|---|
| Agent: planning validity | Workflows were represented as explicit ordered steps with declared inputs, outputs, parameters, and validation rules before local execution; in the frozen held-out evaluation, 11 of 12 supported runs passed every planning, validation, and execution check. | Unlike manual command lines or unconstrained large-language-model code, the plan can be inspected before anything runs. |
| Agent and constraint: frozen held-out evaluation | PaleoRigor passed 23 of 24 held-out runs (95.8%), including all 12 boundary decisions, compared with 19 of 24 (79.2%) for the ablated arm; with a second model configuration the counts were 24 of 24 and 19 of 24. | Workflow success and boundary refusals held for the fixed task set with both model configurations; wide intervals and nonsignificant paired comparisons limit generalization. |
| Reproducibility: public FASTQ source-metadata correspondence | Across six public FASTQ records, PaleoRigor-derived read counts matched ENA/SRA metadata with Δreads = 0, compressed-file ratios were Rbytes = 1, and the MD5 checksums of all 11 files matched ENA's. | The system quantifies correspondence with intended public records at the tested metadata and checksum level before downstream interpretation. |
| Reproducibility: tabular data curation | The peptide validation table was curated from 5,810 rows to 5,696 unique records; 114 duplicate sequence entries were removed. | The result is not only a cleaned table, but a recorded transformation with reusable outputs. |
| Reproducibility: output traceability | Each local run exports three folders: final_outputs (requested results), step_outputs (intermediate files), and ResearchAgent Records (workflow JSON, manifests, checksums, timestamps, and run reports). | The evidence chain can be reviewed later by users, supervisors, or domain experts. |
| Constraint: retraction-associated audit | For SRR29088443, PaleoRigor generated an auditable source-level report while limiting interpretation to file-level reproducibility. | The expert-defined scope kept agreement with repository metadata from being presented as validation of the retracted biological claim. |

This is not a head-to-head speed or biological-accuracy benchmark. The measured-outcome column reports results obtained for PaleoRigor in this study, with ablated-arm counts where applicable. Figure 2 provides a planning-contract ablation using the same model with the PaleoRigor planning contract removed; the remaining comparison statements describe common differences in workflow control and were not measured as external software trials.

### Constraint and reproducibility: reuse of public records from a community catalogue

The three cases above each started from a record chosen for its purpose. To test the system on records as a researcher would take them from a shared resource, we drew 17 ancient-metagenome sequencing records from AncientMetagenomeDir, a community-curated catalogue of ancient metagenomic data [8], using a rule fixed before any file was downloaded: libraries archived at ENA or SRA, with a recorded sample age and 3–50 MB of compressed FASTQ, taking the smallest library of each project (Methods). The records span sample ages from 100 to 52,500 years before present, seven sample materials from dental calculus and teeth to palaeofaeces and a keratinous claw, and Illumina, Ion Torrent, and 454 sequencing. Every catalogued file was downloaded and given to PaleoRigor with the same request, which stated the catalogued library layout and asked for file-type detection, mate pairing for paired-end libraries, FastQC, MultiQC, and a quality-gate report.

Ten of the 17 records (58.8%; Wilson 95% confidence interval, 36.0–78.4%) could be reused exactly as catalogued: they passed all five prespecified checks (Figure 6A,B). Because the records were selected for small file size, this proportion describes these records rather than the catalogue as a whole. No failure was caused by a damaged download. All 29 downloaded files matched their ENA MD5 checksums, the files analyzed in the 15 executed runs had the same Secure Hash Algorithm 256 (SHA-256) checksums as the downloads, and the total base count reported by PaleoRigor’s quality gate equalled ENA’s in all 15. The other seven records failed for four reasons. In two, both the catalogue and ENA listed a paired-end library, but the archive held a single FASTQ file. In one, the archive held both mate files and a third file of 32,854 unpaired reads. In three, ENA’s read count was exactly twice the number of reads in each mate file, because it counted the reads of both mates, whereas for the other seven two-file paired-end records it equalled the number of read pairs. The five paired-end records of the first case all counted the reads of both mates, which is why their read counts were compared after summing both mate files (Figure 3). In one, the catalogued file size and checksum no longer matched the file that ENA now serves (8,523,662 against 3,495,628 bytes), although the read counts agreed.

**Figure 6.**
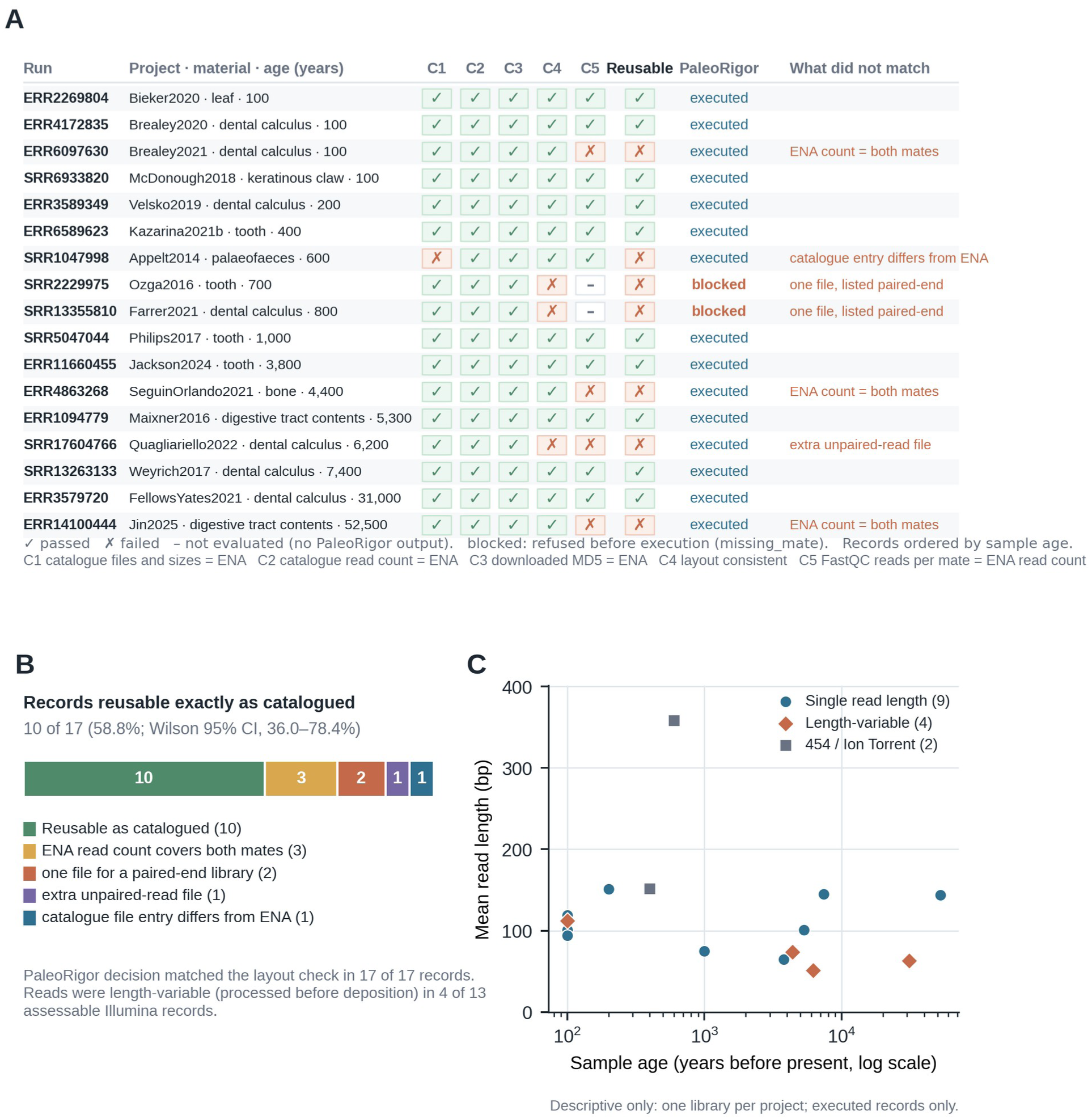
Reuse check of 17 public ancient-metagenome records from AncientMetagenomeDir. A, Prespecified checks for each record, ordered by sample age (years before present): C1, catalogued files, sizes, and MD5 checksums equal those of the current ENA record; C2, the catalogued read count equals ENA’s; C3, the MD5 checksum of every downloaded file equals ENA’s; C4, the catalogued layout equals ENA’s and the file set fits it; C5, the FastQC read count of each mate file equals ENA’s read count. Records passing C1–C5 are reusable as catalogued. The PaleoRigor column shows whether a record was executed or blocked before execution (reason code missing_mate). B, Records reusable as catalogued and the reasons the others failed, with a two-sided Wilson 95% confidence interval. C, Mean read length against sample age for the 15 executed records, by processing state inferred from read lengths; 454 and Ion Torrent reads vary in length by design. The panel is descriptive, and no trend was tested. None of the checks assesses ancient-DNA authenticity.

PaleoRigor’s decision agreed with the file evidence for all 17 records (Figure 6A). It refused both single- file paired-end records before execution with the reason code missing_mate, explaining that mate pairing could not be confirmed without inventing a file. For the record with three files, its plan described the third as an additional FASTQ file, kept it out of the mate pair, and checked it separately; counting this record as a disagreement would give 16 of 17. The five checks themselves were computed afterwards from PaleoRigor’s retained outputs and the ENA records; PaleoRigor decided only whether and how each record was run. Used as catalogued without these checks, two records would have entered a paired-end workflow without a mate file, one would have mixed unpaired reads into a paired analysis, three would have appeared, against ENA’s read count, to have lost half their reads, and one intact download would have failed verification against its catalogued checksum.

The records also differed in what had been done to the reads before deposition. In 4 of the 13 Illumina records that could be assessed (30.8%; 12.7–57.6%), read lengths varied within a file, indicating adapter or quality trimming or read merging before upload; the submitted file name of one of them includes “trimmed”. Such reads pass every agreement check but are not directly comparable with raw reads from another study. Mean read length ranged from 51 to 151 bp in the Illumina records (Figure 6C). Because each project contributed one library, sample age cannot be separated here from laboratory protocol or preprocessing; Figure 6C is descriptive, and no trend was tested.

### Packaged applications for local use

To make the constrained workflow and its records usable outside the development environment, we packaged the system as a self-contained Apple Silicon application for macOS 13 or later that includes seven analysis tools and their runtimes. Analysis of the data files therefore runs on the user’s computer: plans are reviewed in a local interface, validation and the bundled tools run locally, and each run writes its records to a local results folder. The only user data sent to the planning model are the request and uploaded-file summaries. The applications, installation instructions, version information, and SHA-256 checksums are distributed through the project’s GitHub repository [33] and linked from the project website (https://verture-liu.github.io/PaleoRigor/), whose “How it works” page follows a run from request to report. In the release gate for version 0.2.3, recorded on 2 November 2026, all 355 tests in the project test suite passed, and all seven bundled commands returned the expected tool identity or version. The application passed strict ad hoc signature verification and ran from a mounted, read-only disk image. It also removed its temporary launch-token file after use. These results verified packaging on the build and test machine.

A Windows 10/11 x64 installer (version 0.2.3) is produced by a separate native build workflow (GitHub Actions run 34) from the same backend source, and it passed native checks for the installer, the seven bundled tools, backend health and uninstallation. It bundles the same seven analysis tools with an Eclipse Temurin 21.0.8+9 runtime and stores model credentials in Windows Credential Manager; its Samtools build is version 1.24 rather than the 1.23.1 used on macOS. Source URLs and archive checksums for every bundled component are pinned in the repository packaging manifests for both platforms. The Windows build was verified by these native checks only; every result reported in this manuscript was produced on macOS.

## Discussion

PaleoRigor addresses a practical problem that arises before specialist interpretation: researchers can lose track of which files were analyzed and how those files changed. Incorrect inputs, incompatible operations, and undocumented transformations can all produce plausible-looking outputs, and a language-model agent that plans on the researcher’s behalf adds a further risk: a plausible plan that does not fit the data. PaleoRigor answers with three commitments: an agent that states the proposed analysis as an explicit workflow, constraints that bound what that workflow may do, and a retained record from which each run can be reconstructed. Together they make the choice of files, the planned operations, and the resulting changes reviewable before the outputs are used to support biological claims. The evaluation tested each commitment in turn, and the paragraphs below follow the same order.

For the agent and its constraints, the benchmark supports a specific claim. In a post hoc pooled count across the two model configurations, the agent with its planning contract produced a workflow that passed every check in 23 of 24 supported runs (11 of 12 and 12 of 12; the single failure was the .tsv- suffix error, which the validator rejected before execution) and returned the prescribed refusal in all 24 boundary runs. Removing that contract from the same model lowered both counts in both configurations under otherwise identical conditions.

Constraints on interpretation were examined from two directions: the unsupported-claim boundary scenario and the retraction-associated audit. The scenario tests whether the system applies a refusal rule it has been given; the audit shows an expert narrowing the permitted conclusion further than anything the system encodes. Neither is sufficient alone. Encoded rules cannot anticipate an unfamiliar claim, and expert scoping does not scale beyond the analyses an expert personally reviews. Together, the two keep the boundary explicit and reviewable rather than guaranteeing it.

For reproducibility, the retained records did two things. They allowed all 96 run labels from the two frozen model configurations to be recomputed without calling the model again, and every label matched. In the sequencing and table cases, they also showed what traceability adds beyond a run that merely completes: a successful run establishes only that the software did something, whereas the retained records identify which public record it acted on (at the metadata level) and what changed. Both remained recoverable after the fact. That distinction matters most where it is hardest to check by eye, when a researcher decides whether the material is worth interpreting at all. Because a success rate cannot show which record was used or what changed, we present the evidence trail, rather than the success rate, as the part of this system intended to be reused.

For paleobiologists, the practical value is a starting point that can be checked before more demanding analyses proceed. Source and workflow review can precede adapter removal, damage analysis, host removal, taxonomic profiling, contamination assessment, and statistical interpretation [15, 17]. Retaining intermediate evidence allows experts to revisit a choice, identify an error, or reject an analysis while preserving its connection to the original question.

Practical adoption also requires accessible software and inspectable records. The repository retains requests, workflows, intermediate files, checksums, reports, figure sources, and environment details (Additional file 1: Tables S2–S6), and the public website (https://verture-liu.github.io/PaleoRigor/) [27] explains the method and its safeguards. The packaged applications remove the need to install Python, Conda, Java, or the bundled analysis tools separately.

### Limitations

This study evaluates workflow execution, recorded transformations, and agreement with repository metadata; ancient-DNA authentication and biological interpretation were outside its scope. The comparison is an ablation of PaleoRigor’s own planning contract on a fixed task set with one model provider. It was internal by construction, and at 24 runs per arm neither paired comparison reached statistical significance, so the benchmark supports no statement about unfamiliar task types, other providers, or performance relative to established workflow practice. The packaged applications remove installation steps, but whether this improves usability for paleobiologists remains a question for independent user testing.

Broader claims therefore await preregistered evaluations across providers, comparisons with established workflow practice, independent user studies, and evaluations that reflect the uncertainty of archaeological samples more closely. Controlled mixtures could include known amounts of endogenous, environmental, handling, and laboratory material, to assess whether PaleoRigor selects appropriate controls and declines unsupported authenticity claims across contamination levels [34–37]; this would extend evaluation from executing defined tasks to handling uncertainty in ancient biomolecular research. Additional negative controls, such as retracted or disputed studies and low-biomass datasets affected by reagent contamination [35, 37], could show how the system responds to unfamiliar claims, mixed contamination sources, or incomplete archaeological metadata, which neither the present case nor the benchmark establishes.

## Conclusions

Paleomicrobiome studies draw conclusions about past health, diet, and environments from ancient microbial DNA that is scarce, damaged, and prone to contamination, so an error in which files were analyzed, or in how they were changed, can carry through to a biological claim. PaleoRigor addresses this preparatory stage by treating a language-model planner as a bounded research agent. As an agent, it turns a paleobiologist’s research request into an explicit workflow to review before anything runs; in the frozen evaluation, which tested planning, validation, and execution without that review step, 11 of 12 supported held-out runs passed every check. Constraints bound what that workflow may do, including refusing a request to show from quality-control output that reads are ancient or free of contamination: all 12 boundary decisions were correct, and the one planning error among supported runs was rejected by the validator before execution. With its planning contract, PaleoRigor passed 23 of 24 held-out runs in the frozen v5 round, compared with 19 of 24 for the ablated arm, and a preregistered repeat with a second model configuration gave 24 of 24 and 19 of 24. Reproducibility comes from the retained record: all 96 run labels across both model configurations were recomputed from it and matched, six public sequencing records related to ancient or paleomicrobiome research matched repository read counts, compressed-file sizes, and MD5 checksums, and the removal of 114 duplicate sequence entries was documented. For paleomicrobiome research, this means that the inputs, operations, and changes behind a result can be checked before damage analysis, taxonomic profiling, or contamination assessment proceed, while judgments of authenticity and biological meaning stay with the specialist. Within the tested scope, as exercised by its developers, these findings support its use for workflow review and evidence preparation; broader applicability requires the evaluations listed under Limitations. By keeping the agent bounded and its evidence reproducible, PaleoRigor provides a basis for specialist review of fragile ancient microbial evidence.

## Methods

### Agent workflow representation

A user request was represented as an ordered workflow W containing n executable steps:

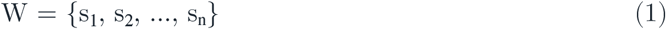

Each step sᵢ was defined by a skill identifier kᵢ, input references Iᵢ, output declarations Oᵢ, parameters θᵢ, and a natural-language reason rᵢ:

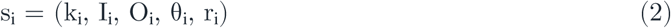

The workflow representation (Equations 1 and 2) records source files, intended outputs, parameters, and reasons before any analysis runs. Each skill identifier refers to a predefined module that can be inspected independently of the planning model. This extends established reproducible-workflow principles with a natural-language planning and review stage [19–21, 38]. Figure 1 relates the representation to Plan, Review, Execute, and Outputs; Table 1 identifies responsibility at each stage.

### Human-in-the-loop checkpoints

Experts review the analysis at three stages. At intake, they check whether the request addresses the scientific question. Before execution, they approve, revise, or reject the proposed workflow and its dependencies. Afterwards, they assess final and intermediate outputs to decide whether further biological analysis is justified. Figure 1 and Additional file 1: Table S1 show the decisions assigned to each checkpoint; the table also lists decisions that arise during local execution and for contested or contamination-sensitive conclusions.

### Workflow validation

The validator, the first constraint on the agent, evaluated a proposed workflow W against the available uploaded files F and the available skill registry K:

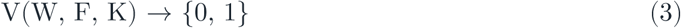

Acceptance required three conditions: each step used a registered skill (kᵢ ∈ K), each input referred to an uploaded file or to an output declared by an earlier step (O<ᵢ denotes the outputs of s₁ … sᵢ₋₁), and input formats matched the skill:

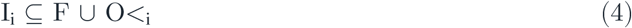

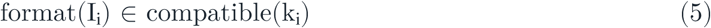

These checks (Equations 3–5; Table 2) identify missing outputs, unresolved references, and incompatible file types before execution. Passing these checks does not establish that a request itself can be answered: it may ask for a conclusion the evidence cannot support, or for an analysis whose required inputs were not supplied or are of the wrong kind. Such requests are handled by a separate constraint, the refusal rules in the planning contract (Engineering benchmark protocol), each returning one of four named reason codes (Results; Additional file 2).

### Execution environment and implemented skills

Execution was the second constraint: validated steps were executed only through predefined local skills. Each skill wrapper connected a workflow step to a tool and recorded inputs, parameters, outputs, warnings, and errors. Table-processing skills standardized peptide comma-separated values (CSV) files, checked sequence syntax, removed duplicate sequences, summarized label counts and peptide lengths, and exported cleaned tables. Sequencing skills supported file detection, pairing checks, FastQC [28], MultiQC [29], SeqKit statistics [30], and source audits. Alignment, subsampling, and host-removal skills wrapped BWA, seqtk, Samtools, and Bowtie 2 [39–42]; none was used in the dataset-level cases. External tools ran in configured environments; missing software was reported to the user without automatic installation.

### Scope and non-goals

PaleoRigor was designed to support workflow review, execution, and documentation. Its checks cover repository metadata, file compatibility, workflow references, quality-control outputs, and run records. Metadata agreement alone cannot establish byte-level identity, ancient-DNA authenticity, contamination sources, microbial ecology, or the validity of a retracted claim. Experts remain responsible for archaeological interpretation. The unsupported-claim boundary scenario of the benchmark tests whether the planner adheres to a predefined limit of this kind, and the retraction- associated audit illustrates the scope of a permitted conclusion (Results).

### Planning-model configuration

All planning calls used the OpenAI-compatible chat-completions endpoint of the model provider, DeepSeek [43]. Each request set the sampling temperature to 0, enabled the provider’s thinking mode, and required a JSON object response, with a 120 s per-request timeout. Transport and Hypertext Transfer Protocol (HTTP) errors were retried up to twice with exponential backoff capped at 4 s. When a returned object failed schema validation, the planner issued exactly one repair call that resupplied the original system prompt together with the validation errors and the invalid object, and instructed the model to correct the JSON without changing the intended task; a second failure aborted the run. In the reported evaluations no request required a retry and no completion required a repair call: all 48 v5 runs and all 48 second-model runs succeeded on the first attempt. The provider exposes no sampling seed, so individual runs are not bit-reproducible.

The frozen v5 evaluation used deepseek-v4-flash and ran on 19 August 2026 between 02:34 and 02:50 Coordinated Universal Time (UTC) at repository commit 4be5f2f. The preregistered second-model check used deepseek-v4-pro and ran on 19 August 2026 between 06:52 and 07:14 UTC at commit 728a226.

The only change to the evaluation code between those commits was the addition of per-model configuration loading; the scenarios, prompts, schemas, scoring rules, and agent source were unchanged. Both commits are contained in release v0.2.0 (commit 4febac7), the planning prompts were not changed between either commit and the release, and the benchmark code was not changed between commit 728a226 and the release. Each run stores a provenance record containing the arm, start and end times, the model string reported by the interface, token usage, latency, attempt count, repair-call count, and the repository commit. The complete system prompts for both arms are reproduced in Additional file 2.

### Engineering benchmark protocol

The benchmark tested the agent and its constraints together in three versioned rounds, v3, v4, and v5, each comparing a PaleoRigor arm with an ablated arm in a fixed paired design with eight scenarios: four supported and four boundary scenarios. Three repetitions per scenario in each arm produced 24 runs per arm and 48 application programming interface (API) calls per round. The arms used the same model configuration, with calls made in alternating order. Both arms received the same workflow schema, blocked-decision schema, uploaded-file summaries, and skill registry; this shared block was byte- identical at 35,682 characters. The ablated arm received, in addition, only a four-sentence planner instruction of 247 characters. The PaleoRigor arm instead received the full planning contract of 40,347 characters, comprising 30 rules governing workflow stages, file compatibility, prerequisites, and declared outputs, one worked example workflow, and six refusal rules that state the four boundary reason codes together with the limits on scientific claims. Supported workflows were executed locally by the harness directly after validation, without the interactive approval step, so the benchmark tested planning, validation, and execution rather than expert review. Boundary requests were evaluated as decisions and were never executed.

PaleoRigor passed 18 of 24 runs in each of v3 and v4, which were retained as failed qualification rounds. Their failures informed development and were not reclassified as confirmatory successes. Before v5, we corrected output-name mapping, FASTA/FASTQ suffix handling, peptide-statistics declarations, chart aliases, and metadata-only sample-sheet planning. We froze the v5 manifest, inputs, prompts, scoring rules, threshold, and code before the first API call. The v5 round used new files, identifiers, values, and wording, with a fixed sample size and no selective reruns or replacement of failures.

For each run r and each criterion d in its required set Dᵣ, qᵣ,d was 1 when the run passed the stated rule and 0 when it failed. Supported-run criteria covered parsing, required and forbidden skills, production workflow validation, and local execution. Boundary runs required a valid blocked response with the prespecified reason code. A run passed only when every relevant rule passed:

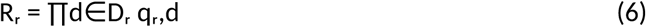

For scenario t with nₜ repeated runs, the pass rate was calculated as:

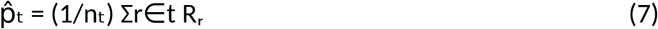

Strict-success proportions, from the run-level outcomes of Equation 6, are reported with two-sided Wilson 95% confidence intervals. The prespecified v5 criterion required at least 22 of 24 PaleoRigor runs (more than 90%) to pass. Paired differences are reported in percentage points and assessed with an exact two-sided McNemar test; this secondary comparison had limited power with 24 pairs, which comprise eight scenarios each repeated three times with identical inputs and are therefore not fully independent. Each v5 label was recomputed from its retained response, workflow or blocked decision, validation result, and execution record. All 48 labels matched. Scenario-level pass rates (Equation 7) are shown in Figure 2C as successes out of three. Manifests, scores, summaries, and Figure 2 source data are retained under analysis/benchmark_v3, analysis/benchmark_v4, analysis/benchmark_v5, and figures/manuscript/benchmark_v5; the second-model check and the incomplete third configuration are retained under analysis/benchmark_multimodel.

For the second-model check, we reused the frozen v5 design and changed the requested model from deepseek-v4-flash to deepseek-v4-pro. The eight scenarios, three repetitions, two arms, prompts, temperature (0), thinking mode, and JSON response mode were unchanged. Validation, scoring, local execution, and the fixed denominator were also retained. The second-model hypothesis and decision rule were committed before the first formal call. The criterion required at least 22 of 24 PaleoRigor successes and more successes than the ablated arm. Recomputing all 48 second-model labels from retained records confirmed every label; no API call failed. The previously observed deepseek-v4-flash results provided descriptive context only.

A third model configuration, deepseek-v4-flash-vision-exp, was also run against the frozen v5 design on 21 August 2026, but the experiment was not completed: only the ablated arm was executed (17 of 24 successes, 70.8%; Wilson 95% confidence interval, 50.8–85.1%). No matched PaleoRigor arm was run, so this configuration yields no paired comparison and contributes no result to the comparisons reported here. Its manifest, run records, and summary are retained in the repository and are reported here so that the retained evaluation materials can be reconciled with the results presented above.

### Dataset-level evaluations and rigor criteria

The evaluation comprised three versioned benchmark rounds, a preregistered second-model check, and three dataset-level case studies (Table 3). Rounds v3 and v4 informed development, whereas v5 evaluated the frozen system on held-out inputs. Two cases addressed reproducibility: the sequencing source checks of six public records tested agreement with public metadata (Figure 3), and the peptide- table curation tested whether data changes were recorded (Figure 4). The retraction-associated audit examined an expert-defined limit on interpretation, the constraint seen from the expert’s side (Figure 5). Outcomes included strict workflow success, correct boundary decisions, duplicate removal, table structure, read counts, compressed-file sizes, and descriptive quality-control measures.

### Source-level agreement metrics

Read-count and compressed-file-size agreement were calculated against public ENA/SRA metadata:

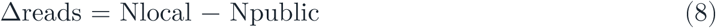

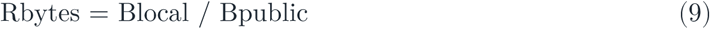

Agreement at the measured metadata level required Δreads = 0 and Rbytes = 1 (Equations 8 and 9). Here, Δreads denotes the difference between local and public read counts; Rbytes is the ratio of local compressed FASTQ size to public fastq_bytes. Run records retained accessions, library layout, and mate organization. R1 and R2 denote paired-end mate files. Figure 3 applies these measures to the sequencing source checks, and Figure 5B compares the same quantities for the retraction-associated audit.

Δreads and Rbytes alone cannot exclude files that differ in content but have identical read counts and compressed sizes, so content identity was checked separately with checksums. The FASTQ files analysed in the sequencing source checks and the retraction-associated audit are kept in the repository. For each of these 13 files, the SHA-256 checksum that PaleoRigor recorded for the uploaded input in its run record equalled that of the kept copy, and the MD5 checksum of the kept copy equalled the fastq_md5 value published by the ENA Portal API (retrieved on 24 October 2026). The comparison script and its output are in analysis/source_identity.

### FASTQ quality-control summaries

FASTQ summaries included GC percentage, mean read length, and the fraction of bases recorded as N. For a FASTQ file with m reads and read lengths Lⱼ, the measures were:

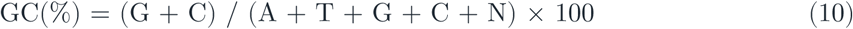

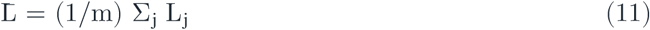

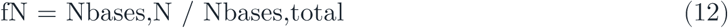

FastQC and MultiQC generated standard quality-control files [28, 29]. SeqKit produced short file-level summaries used in the figures [30]. GC content, mean read length, and N-fraction (Equations 10–12) are reported descriptively in Figure 5C; GC content is also shown for the sequencing source checks in Figure 3C.

### Output organization, public website, and packaged desktop applications

Each local run created a dated result folder. final_outputs contained the files requested by the user. step_outputs contained intermediate files. ResearchAgent Records (the run-record folder, named by the application) contained workflows, manifests, checksums, timestamps, and reports. This structure keeps both the requested result and the evidence used to produce it, so that each run can be checked from its record. The public website (https://verture-liu.github.io/PaleoRigor/) [27] explains the method and installation process; it does not accept research data or run workflows. Analyses of user data run locally in the browser interface, where users review the workflow, approve execution, monitor progress, and inspect the report. On macOS, this interface is launched by the packaged desktop application, a self- contained Apple Silicon application targeting macOS 13 or later. The application bundles the PaleoRigor backend together with FastQC 0.12.1 [28], MultiQC 1.35 [29], SeqKit 2.13.0 [30], seqtk 1.5-r133 [39],

Samtools 1.23.1 [40], BWA 0.7.19-r1273 [41], Bowtie 2 (version 2.5.5) [42], and their required runtimes. Users therefore do not need to install Python, Conda, Java, or these seven tools separately. The backend binds only to the local loopback interface, uses a temporary per-launch authentication token, and stores model API credentials in macOS Keychain rather than in the application bundle. The Windows installer exposes the same local browser interface from the same backend source (Results).

### Reproducibility bundle

Each case retained final outputs, intermediate files, and run records. The public FASTQ files analysed in the dataset-level cases are kept in the repository with their accessions and ENA metadata. The repository contains processed outputs, workflows, quality-control summaries, manifests, checksums, and plotting scripts. It also includes the v3–v5 benchmark manifests, the preregistered second-model check, run-level scores, summaries, verification reports, and Figure 2 source data. In Additional file 1, Tables S2–S4 link the reported results to these records, Table S5 links the errors observed during development to the checks that now prevent them, and Table S6 records the software environment. The plotting scripts use Python 3.13.9 [44], pandas 2.3.3 [45], and Matplotlib 3.11.0 [46].

The evaluation was designed to be reconstructible rather than merely described. The hypothesis, decision rule, scenario manifest, input files, prompts, scoring rules, and success threshold were committed before the first formal API call of each round, and the preregistrations are kept in the repository. The scenarios themselves were held out: the frozen v5 round used files, identifiers, values, and request wording that had not been seen during development, and no run was rerun or replaced after the fact. Development-stage qualification rounds and their failures were retained rather than removed. Every run writes the provenance record described under Planning-model configuration, alongside the verbatim request, the returned plan or blocked decision, the validation result, and the execution record. From those records, all 96 run labels across the two model configurations were recomputed, and all 96 matched. The software environment is pinned, the analysis outputs carry checksums, and the releases corresponding to these results are archived under persistent identifiers (Availability of data and materials). A reader who disagrees with a reported number can therefore recompute it from the deposited records without rerunning the language model, whose outputs are not bit-reproducible.

### Public-data reuse check

For the reuse check (Figure 6), we used the host-associated metagenome sample and library tables of AncientMetagenomeDir [8] at commit 0a39020 (16 October 2026), which list 2,980 libraries. The selection rule, request texts, checks, and outcome measures were written into a protocol before any file was downloaded. Libraries archived at ENA or SRA with download links, a recorded sample age, and 3– 50 MB of catalogued FASTQ were kept, and the smallest library of each project was chosen, giving 17 records. On 19 October 2026, we retrieved the current ENA read-run report of each record [7] and downloaded every catalogued FASTQ file without renaming, decompressing, or otherwise changing it (29 files, 199.8 MB). Each record was then run as a separate task in the PaleoRigor application with one of two fixed requests, which differed only in the catalogued library layout; the author approved each valid plan without editing it, and refusals, which export no files, were kept as screenshots. Five checks (C1–C5; Figure 6 legend) were computed from the catalogue, the ENA reports, the download log, and PaleoRigor’s exported outputs. The primary outcome was the proportion of records passing all five, reported with a two-sided Wilson 95% confidence interval. Secondary outcomes were PaleoRigor’s decision for each record and its agreement with C4 and, for Illumina records, whether the reads within a file had a single length or several. The protocol, the selection, download, summary, and plotting scripts, the request texts, and the source data are in analysis/public_reuse_check; the catalogue copies, ENA reports, and download log are in data/raw/public_reuse_check; and per-record results are listed in Additional file 1: Table S7.

### Use of AI-assisted tools

Large-language-model assistants were used during software development and manuscript preparation, to draft and refactor code and to edit the language and structure of the text. All generated code was reviewed and tested by the authors, and all analyses, results, and conclusions were checked by the authors, who take full responsibility for the content of this article. These assistants are distinct from the planning models used inside PaleoRigor and in its evaluation, which are described under Planning-model configuration.

## List of abbreviations

API: application programming interface
bp: base pairs
CSV: comma-separated values
DOI: digital object identifier
ENA: European Nucleotide Archive
GC: guanine–cytosine
HTTP: Hypertext Transfer Protocol
JSON: JavaScript Object Notation
LLM: large language model
MD5: Message-Digest algorithm 5
QC: quality control
SHA-256: Secure Hash Algorithm 256
SRA: Sequence Read Archive
UTC: Coordinated Universal Time.

## Declarations

### Ethics approval and consent to participate

Not applicable. This study performed secondary computational analyses of public repository records and small example files and did not recruit participants, collect new human or animal material, or generate new human-subject data.

### Consent for publication

Not applicable.

### Availability of data and materials

The datasets analyzed in this study are public repository records or small example files. Public sequence inputs are identified by accession in Table 3. Evaluation manifests, run-level scores, workflows, blocked decisions, execution records, processed outputs, figure source data, and plotting scripts are available in the PaleoRigor repository [33]. The public FASTQ files analysed in this study are also kept in the repository (stored with Git Large File Storage), and their accession identifiers enable retrieval from ENA or SRA [6, 7].

Project name: PaleoRigor. Project home page: https://verture-liu.github.io/PaleoRigor/ [27]. Source code: https://github.com/Verture-Liu/PaleoRigor [33]. Desktop applications, installation instructions, version information, and SHA-256 checksums: https://github.com/Verture-Liu/PaleoRigor/tree/main/paleorigor. Archived versions: the current release, v0.2.3 (commit 506cfdd), is archived at Zenodo (https://doi.org/10.5281/zenodo.23883541). The reported results were produced with code contained in release tag v0.2.0 (commit 4febac7), archived at Zenodo (https://doi.org/10.5281/zenodo.22912729). Release v0.2.2 (commit a7bca3b), archived at https://doi.org/10.5281/zenodo.23042955, adds the per-run records of the benchmark rounds without modifying the earlier benchmark files; the concept digital object identifier (DOI) https://doi.org/10.5281/zenodo.22912728 always resolves to the latest version. Folder paths cited in this article and its additional files follow the current repository layout; releases up to and including v0.2.2 use earlier folder names, which results/README.md in the repository maps to the current ones. Operating systems: Apple Silicon macOS 13 or later, and Windows 10/11 x64 (packaged applications). Linux is not supported in this release: no Linux package is provided, and the source version stores model credentials only in macOS Keychain or Windows Credential Manager. All reported results were produced on macOS; the Windows build was verified by its native build, installer, tool-identity, backend and uninstall checks only and produced no reported result. Programming language: Python 3.13.9. License:

MIT. Redistributed third-party tools retain their own licenses, several of which are GPL-3.0; these are listed with their source obligations in THIRD_PARTY_NOTICES.md in the repository. Restrictions on non- academic use: none.

### Authors’ contributions

TL designed and implemented PaleoRigor, performed the evaluations and analyses, and wrote the manuscript. HZ contributed ideas to the project and advised on key design decisions. YJ tested the software and revised the manuscript. KN supervised the project. All authors read and approved the final manuscript.

## Acknowledgements

Not applicable.

## Additional files

Additional file 1 (.pdf). Supplementary tables and note. Tables S1–S7 (expert checkpoints; links to final outputs, intermediate files, and run records; evidence files; claim-to-evidence links; errors and implemented controls; software and environment; per-record results of the public-data reuse check) and a supplementary note on figure generation and image provenance.

Additional file 2 (.zip). System prompts. The verbatim system prompts sent to both benchmark arms, with the byte-identical shared block and the content unique to each arm.

