## Supplementary Information for "PaleoRigor: a bounded research agent with a reproducible evidence trail for paleomicrobiome data processing"

### Additional file 1

Reference numbers in square brackets refer to the reference list of the main manuscript.

This file contains seven tables and a supplementary note. Table S1 sets out the expert checkpoints. Table S2 links each case to final outputs, intermediate files, and run records, and Tables S3–S6 cover case evidence, claim-to-evidence links, error controls, and software settings. Table S7 lists the per-record results of the public-data reuse check. Additional file 2 contains the verbatim system prompts sent to both benchmark arms, together with the byte-identical shared block and the content unique to each arm. The workflows, reports, manifests, and checksums of each run, the example final_outputs and step_outputs, and the figure source data and plotting scripts are not duplicated here: they are available in the archived PaleoRigor repository (Availability of data and materials in the main manuscript), in the result folders listed in Table S2, under figures/manuscript, and in the figure scripts under analysis. Together, these materials support Figures 1–6 and Tables 1–4.

**Table S1. Expert checkpoints. The table shows where expert review occurs and which error each checkpoint is intended to prevent.**

| Workflow stage | PaleoRigor action | Expert decision | Failure prevented |
|---|---|---|---|
| Task intake | Converts natural-language request into structured workflow steps with declared inputs, outputs, and parameters. | Check whether the planned task matches the scientific question before execution. | Wrong task framing or use of an irrelevant skill. |
| Pre-execution validation | Checks file references, step-output dependencies, and input-format compatibility. | Approve, revise, or reject the proposed workflow. | Hallucinated step references, CSV/FASTQ mismatch, or unsupported operation. |
| Local execution | Runs only predefined skills and reports progress, warnings, and missing environment dependencies. | Decide whether to install missing tools or stop the run. | Silent auto-installation, uncontrolled code execution, or hidden dependency failure. |
| Post-run review | Exports final_outputs, step_outputs, and ResearchAgent Records. | Interpret outputs and decide whether downstream biological analysis is justified. | Treating quality-control (QC) or source checks as biological validation. |
| Boundary cases | Flags source-level reproducibility separately from claim-level interpretation. | Gate contested or contamination-sensitive conclusions for domain review. | Overclaiming from low-biomass or retraction-associated records. |

**Table S2. Links to final outputs, intermediate files, and run records. final_outputs contains requested result files. step_outputs contains files produced by each workflow step. ResearchAgent Records contains workflow JSON, manifests, timestamps, checksums, and reports. Each row links a case or sequencing record to the repository folder used to review its evidence trail; ERR15682270 has two rows because its two mates were processed in separate runs. Paths follow the current repository layout; results/README.md in the repository maps them to the folder names used in releases up to and including v0.2.2.**

| Validation item | Result folder | Repository location | Evidence role |
| --- | --- | --- | --- |
| Case 1a: ERR15682270 R1 | results/ sequencing_source_checks/ ERR15682270_R1 | PaleoRigor repository path: results/sequencing_source_checks/ERR15682270_R1 [33] | FASTQ file typing, FastQC/MultiQC outputs, quality-gate report, and run records for mate R1. Subfolders at this repository location contain final_outputs, step_outputs, and ResearchAgent Records. |
| Case 1b: ERR15682270 R2 | results/ sequencing_source_checks/ ERR15682270_R2 | PaleoRigor repository path: results/sequencing_source_checks/ERR15682270_R2 [33] | FASTQ file typing, FastQC/MultiQC outputs, quality-gate report, and run records for mate R2. Subfolders at this repository location contain final_outputs, step_outputs, and ResearchAgent Records. |
| Case 1c: ERR10114877 paired FASTQ | results/ sequencing_source_checks/ ERR10114877 | PaleoRigor repository path: results/sequencing_source_checks/ERR10114877 [33] | Paired-end FASTQ QC outputs and source-record audit records after a rerun with corrected inputs. Earlier runs that processed each mate separately (results/earlier_runs/ERR10114877_R1_single_file and ERR10114877_R2_single_file) give the same read counts and GC content and supply this record's Figure 3 source data. Subfolders at this repository location contain final_outputs, step_outputs, and ResearchAgent Records. |
| Case 1d: ERR3250149 single-end FASTQ | results/ sequencing_source_checks/ ERR3250149 | PaleoRigor repository path: results/sequencing_source_checks/ERR3250149 [33] | Single-end FASTQ QC outputs, source-record checks, and local run records. Subfolders at this repository location contain final_outputs, step_outputs, and ResearchAgent Records. |
| Case 1e: ERR10114867 paired FASTQ | results/ sequencing_source_checks/ ERR10114867 | PaleoRigor repository path: results/sequencing_source_checks/ERR10114867 [33] | Paired-end FASTQ QC outputs and source-record audit records after a rerun with corrected inputs. An earlier run (results/earlier_runs/ERR10114867_first_run) gives the same read counts and GC content and supplies this record's Figure 3 source data. Subfolders at this repository location contain final_outputs, step_outputs, and ResearchAgent Records. |

| Validation item | Result folder | Repository location | Evidence role |
|---|---|---|---|
| Case 1f: ERR10114861 paired FASTQ | results/ sequencing_source_checks/ ERR10114861 | PaleoRigor repository path: results/sequencing_source_checks/ERR10114861 [33] | Paired-end FASTQ QC outputs, metadata checks, and run records. Subfolders at this repository location contain final_outputs, step_outputs, and ResearchAgent Records. |
| Case 1g: ERR15682267 paired FASTQ | results/ sequencing_source_checks/ ERR15682267 | PaleoRigor repository path: results/sequencing_source_checks/ERR15682267 [33] | Paired-end FASTQ QC outputs, metadata checks, and run records. Subfolders at this repository location contain final_outputs, step_outputs, and ResearchAgent Records. |
| Case 2: peptide-table curation | results/peptide_table_curation | PaleoRigor repository path: results/peptide_table_curation [33] | Cleaned CSV, peptide statistics, charts, normalization/validation/deduplication step outputs, workflow manifest, and report. Subfolders at this repository location contain final_outputs, step_outputs, and ResearchAgent Records. |
| Case 3: retraction-associated audit | results/ retraction_associated_audit | PaleoRigor repository path: results/retraction_associated_audit [33] | Boundary-aware source-data audit, SeqKit/FastQC/MultiQC summaries, intermediate step outputs, and run records. Subfolders at this repository location contain final_outputs, step_outputs, and ResearchAgent Records. |
| Engineering benchmark | analysis/benchmark_v3; analysis/benchmark_v4; analysis/benchmark_v5; analysis/benchmark_multimodel; figures/manuscript/benchmark_v5 | PaleoRigor repository paths: analysis/benchmark_v3, analysis/benchmark_v4, analysis/benchmark_v5, analysis/benchmark_multimodel, and figures/manuscript/benchmark_v5 [33] | Versioned manifests, preregistrations, run-level scores, summary JSON, verification report, and Figure 2 source data. The runs subfolder of each round contains, for every run, the request, raw completion, workflow or blocked decision, score, and provenance record. |

**Table S3. Evidence files for each case. The table links each case to its natural-language request, workflow records, final outputs, and intermediate files.**

| Case | Task and workflow evidence | Final-output evidence | Intermediate evidence |
|---|---|---|---|
| Case 1: sequencing source checks | FASTQ source-check and QC requests for ERR15682270, ERR10114877, ERR3250149, ERR10114867, ERR10114861, and ERR15682267. | FastQC/MultiQC summaries and metadata JSON files; repository paths listed in Table S2. | Per-record step_outputs and ResearchAgent Records; repository paths listed in Table S2. |
| Case 2: peptide-table curation | Natural-language peptide-table curation request; workflow JSON and report in ResearchAgent Records. | Cleaned CSV, statistics, and charts; repository path listed in Table S2. | Normalization, validation, statistics, deduplication, and export step outputs; repository path listed in Table S2. |
| Case 3: retraction-associated audit | Boundary-aware audit request for SRR29088443; workflow report documents that source-data reproducibility rather than biological validity was tested. | Audit summaries and QC outputs; repository path listed in Table S2. | File detection, FastQC, MultiQC, SeqKit, and run records; repository path listed in Table S2. |
| Engineering benchmark | Benchmark requests for rounds v3–v5 and the second-model check; versioned manifests, preregistrations, raw completions, workflows or blocked decisions, and validation and execution records. | v5 run_level.csv, summary.json, verification_report.md, independent scoring outputs, Figure 2 source data, and SVG/PDF/PNG/TIFF exports; repository paths listed in Table S2. | Per-run records for v3–v5 under analysis/benchmark_v3, analysis/benchmark_v4, and analysis/benchmark_v5, Figure 2 source data under figures/manuscript/benchmark_v5, and second-model records under analysis/benchmark_multimodel; repository paths listed in Table S2. |

**Table S4. Links between manuscript claims and evidence. Each claim is paired with the file or record that supports it.**

| Manuscript claim | Primary evidence file or record | Figure/Table location | Repository route |
|---|---|---|---|
| Public FASTQ source records matched local inputs at the read-count and compressed-file-size level. | fastqc_run_metadata.json, file_type_report.json, MultiQC output, and run records for Cases 1a–1g. | Figure 3; Table 3 and Table S2 | Case 1a–1g repository paths in Table S2 |
| Peptide-table curation removed 114 duplicate sequence entries and exported 5,696 unique records. | cleaned_Validation.csv, peptide_statistics.json, and deduplication step outputs. | Figure 4; Table 3 | Case 2 repository path in Table S2 |
| The retraction-associated audit reproduced source-level metadata without validating the retracted biological conclusion. | seqkit_stats.tsv, fastqc_run_metadata.json, workflow report, and boundary-aware audit figure source data. | Figure 5; Table 3 and Tables S2–S3 | Case 3 repository path in Table S2 |
| The system separates user-facing final outputs from intermediate step outputs and run records. | Timestamped result folders containing final_outputs, step_outputs, and ResearchAgent Records. | Table S2 | Repository result folders listed in Table S2 |
| Workflow validation prevents unsupported execution such as missing step outputs or incompatible file-type operations. | Workflow validation errors and development failure cases documented in run records and Table S5. | Table 2 and Table S5 | ResearchAgent Records under the repository paths in Table S2 |
| The frozen v5 round exceeded the prespecified strict-success threshold, while earlier qualification failures remained visible. | v3–v5 manifests and summaries; v5 run_level.csv, summary.json, verification_report.md, and retained execution records. | Figure 2; Table 4 | Engineering benchmark folders in Table S2 (analysis/benchmark_v3, analysis/benchmark_v4, analysis/benchmark_v5, analysis/benchmark_multimodel, and the Figure 2 source-data folder) |

**Table S5. Errors and implemented controls. Development failures were kept because they show why each safety check was needed.**

| Observed failure mode | How it appeared | Implemented control | Evidence route |
|---|---|---|---|
| Missing step-output reference | Generated workflow referred to outputs such as step_02:qc_zip or step_01/normalized that the validator could not resolve. | Strict step-output reference validation before execution. | Workflow validation run records |
| Wrong input type for requested operation | FASTQ QC was requested for a CSV upload, or gzip decompression was proposed for an already FASTQ-classified file. | File-type detection and skill compatibility checks block incompatible workflows. | Workflow records and file-type compatibility checks |
| Quality gate stopping on expected duplicates | The peptide validation file contained duplicate sequences, causing a strict gate to fail before deduplication. | Deduplication and quality-gate order clarified; warnings can be audited before downstream use. | Case 2 step_outputs and deduplication records |
| Tool-output directory assumption | FastQC initially failed when its output directory was not created before execution. | Skill wrapper creates required output directories and records tool metadata. | FASTQ final_outputs and tool metadata |
| User prompt/file-name mismatch | A rerun failed when the natural-language prompt still named files from a previous group. | Workflow review exposes uploaded-file references before local execution. | Corrected rerun records |

**Table S6. Software and environment details. The table lists the local software used to generate the evaluation outputs and figures. The archived release corresponding to these results is identified under Availability of data and materials in the main manuscript.**

| Component | Version, environment, and source | Role in study | Release note |
|---|---|---|---|
| PaleoRigor project | Repository: PaleoRigor GitHub repository [33]; release tag v0.2.0, commit 4febac7 | Workflow planning, validation, local skill execution, and output organization. | Archived as tagged GitHub releases with Zenodo DOIs 10.5281/zenodo.22912729 (v0.2.0), 10.5281/zenodo.23042955 (v0.2.2) and 10.5281/zenodo.23883541 (v0.2.3). |
| Python runtime for manuscript scripts | Python 3.13.9 in project virtual environment; Python [44]; pandas [45]; Matplotlib [46] | Figure source-data processing and manuscript figure generation. | Used with pandas 2.3.3 and Matplotlib 3.11.0. |
| FastQC | FastQC 0.12.1; software reference [28] | FASTQ quality-control reports for the sequencing source checks and the retraction-associated audit. | Called through the predefined fastq_qc skill; reports preserved in final_outputs. |
| MultiQC | MultiQC 1.35 in multiqc_env; software reference [29] | Aggregation of FastQC reports into summary HTML/metadata outputs. | Missing environment is reported rather than silently installed. |
| SeqKit | SeqKit 2.13.0 in base environment; software reference [30] | Compact FASTQ statistics and source-level summaries. | Used for read/base/length checks where available. |
| seqtk | seqtk 1.5-r133, bundled with the packaged applications; software reference [39] | Read subsampling skill (seqtk_sample). | Bundled tool identity checked in the release gate; not used by the three dataset-level cases. |
| Samtools | Samtools 1.23.1 (macOS) and 1.24 (Windows), bundled with the packaged applications; software reference [40] | Alignment statistics, sorting, and indexing skills (samtools_stats, samtools_sort_index). | Bundled tool identity checked in the release gate; not used by the three dataset-level cases. |
| BWA | BWA 0.7.19-r1273, bundled with the packaged applications; software reference [41] | Read-alignment skill (bwa_align). | Bundled tool identity checked in the release gate; not used by the three dataset-level cases. |
| Bowtie 2 | Bowtie 2 (version 2.5.5), bundled with the packaged applications; software reference [42] | Host-read removal skill (host_dna_removal). | Bundled tool identity checked in the release gate; not used by the three dataset-level cases. |
| Planning models | deepseek-v4-flash (frozen v5 evaluation) and deepseek-v4-pro (second-model check) through the provider's chat-completions endpoint; provider documentation [43] | Workflow planning in both benchmark arms. | Temperature 0, thinking mode, and JSON object responses; the provider exposes no sampling seed. |

| Component | Version, environment, and source | Role in study | Release note |
| --- | --- | --- | --- |
| Repository and archives | GitHub repository: PaleoRigor GitHub repository [33]; archived releases: Zenodo DOIs 10.5281/zenodo.22912729 (v0.2.0), 10.5281/zenodo.23042955 (v0.2.2) and 10.5281/zenodo.23883541 (v0.2.3) | Stable access to code, outputs, step files, run records, and figure source data. | Reported results correspond to release v0.2.0; release v0.2.2 adds the per-run benchmark records, and release v0.2.3 (10.5281/zenodo.23883541) adds the source-identity and public-data reuse checks; the evaluation code and reported results are unchanged. Cite the archived snapshots rather than the mutable main branch. |

**Table S7. Per-record results of the public-data reuse check. Records are ordered by sample age (years before present). Checks C1–C5 are defined in the legend of Figure 6 in the main manuscript; a record is reusable as catalogued when it passes all five. Read lengths are the range across the record's files in PaleoRigor's quality-gate report; 454 and Ion Torrent reads vary in length by design. The source data are in analysis/public_reuse_check/source_data/case4_records.csv and case4_files.csv.**

| Run | Project; material; age (years) | Instrument; catalogued layout | ENA read count | Checks not passed | PaleoRigor | Read lengths |
|---|---|---|---|---|---|---|
| ERR2269804 | Bieker2020; leaf; 100 | Illumina HiSeq 2000; paired-end, 2 files | 52,276 | None | Executed | 101 bp, single length |
| ERR4172835 | Brealey2020; dental calculus; 100 | Illumina HiSeq 2500; paired-end, 2 files | 38,940 | None | Executed | 119 bp, single length |
| ERR6097630 | Brealey2021; dental calculus; 100 | Illumina NovaSeq 6000; paired-end, 2 files | 165,028 | C5: ENA read count covers both mates | Executed | 94 bp, single length |
| SRR6933820 | McDonough2018; keratinous claw; 100 | Illumina MiSeq; paired-end, 2 files | 235,197 | None | Executed | 35–151 bp, variable |
| ERR3589349 | Velsko2019; dental calculus; 200 | Illumina HiSeq 2500; paired-end, 2 files | 28,517 | None | Executed | 151 bp, single length |
| ERR6589623 | Kazarina2021b; tooth; 400 | Ion Torrent Proton; single-end, 1 file | 195,972 | None | Executed | 25–345 bp (not assessable) |
| SRR1047998 | Appelt2014; palaeofaeces; 600 | 454 GS FLX Titanium; single-end, 1 file | 29,811 | C1: catalogued size and MD5 differ from ENA | Executed | 50–574 bp (not assessable) |
| SRR2229975 | Ozga2016; tooth; 700 | Illumina HiSeq 2500; paired-end, 1 file | 132,145 | C4, C5: one file for a paired-end library | Blocked (missing_mate) | Not run |
| SRR13355810 | Farrer2021; dental calculus; 800 | Illumina HiSeq 2500; paired-end, 1 file | 140,430 | C4, C5: one file for a paired-end library | Blocked (missing_mate) | Not run |
| SRR5047044 | Philips2017; tooth; 1,000 | Illumina Genome Analyzer II; single-end, 1 file | 101,826 | None | Executed | 75 bp, single length |
| ERR11660455 | Jackson2024; tooth; 3,800 | Illumina MiSeq; single-end, 1 file | 168,845 | None | Executed | 65 bp, single length |
| ERR4863268 | SeguinOrlando2021; bone; 4,400 | Illumina MiniSeq; paired-end, 2 files | 105,478 | C5: ENA read count covers both mates | Executed | 67–74 bp, variable |
| ERR1094779 | Maixner2016; digestive tract contents; 5,300 | Illumina HiSeq 2000; paired-end, 2 files | 237,328 | None | Executed | 101 bp, single length |
| SRR17604766 | Quagliariello2022; dental calculus; 6,200 | Illumina NovaSeq 6000; paired-end, 3 files | 318,602 | C4, C5: extra file of unpaired reads | Executed | 27–51 bp, variable |
| SRR13263133 | Weyrich2017; dental calculus; 7,400 | Illumina HiSeq 2500; paired-end, 2 files | 53,145 | None | Executed | 145 bp, single length |
| ERR3579720 | FellowsYates2021; dental calculus; 31,000 | NextSeq 500; paired-end, 2 files | 79,655 | None | Executed | 1–76 bp, variable |
| ERR14100444 | Jin2025; digestive tract contents; 52,500 | Illumina NovaSeq 6000; paired-end, 2 files | 242,902 | C5: ENA read count covers both mates | Executed | 144 bp, single length |

### Supplementary note: figure generation and image provenance

All figures were produced from author-created vector schematics or Python-based plots of analytical outputs. The graphical abstract contains no photographs; its excavation scene is an author-drawn vector schematic. No figure was generated with a text-to-image model. Figure 2 was generated in Python from the retained v3–v5 benchmark summaries and run-level records. Its source-data CSV, editable SVG and PDF exports, and PNG and TIFF exports are in the repository under figures/manuscript/benchmark_v5, and its plotting script is analysis/benchmark_v5/plot_v5_benchmark.py. Figure 1 and the graphical abstract are drawn as SVG by the scripts in analysis/schematics, Figures 3–5 by analysis/make_task_grouped_figures.py and analysis/make_retracted_case_figure_v2.py, and Figure 6 by analysis/public_reuse_check/plot_case4.py from the source data in analysis/public_reuse_check/source_data.
